# Psychophysical Scaling Reveals a Unified Theory of Visual Memory Strength

**DOI:** 10.1101/325472

**Authors:** Mark W. Schurgin, John T. Wixted, Timothy F. Brady

## Abstract

Almost all models of visual memory implicitly assume that errors in mnemonic representations are linearly related to distance in stimulus space. Here, we show that neither memory nor perception are appropriately scaled in stimulus space; instead, they are based on a transformed similarity representation that is non-linearly related to stimulus space. This result calls into question a foundational assumption of extant models of visual working memory. Once psychophysical similarity is taken into account, aspects of memory that have been thought to demonstrate a fixed working memory capacity of ~3-4 items and to require fundamentally different representations -- across different stimuli, tasks, and types of memory -- can be parsimoniously explained with a unitary signal detection framework. These results have significant implications for the study of visual memory and lead to a substantial reinterpretation of the relationship between perception, working memory and long-term memory.

---

Working memory is typically conceptualized as a fixed capacity system, with a discrete number of items, each represented with a certain degree of precision^1,2^. It is thought to be a core cognitive system^3,4^, with individual capacity differences strongly correlating with measures of broad cognitive function such as fluid intelligence and academic performance^5,6^. As a result, many researchers are deeply interested in understanding and quantifying working memory capacity and understanding the connections between working memory and long-term memory.

Continuous feature spaces are often used to investigate memory, as they allow the precise quantification of information stored in memory^2,7,8^. In one prominent paradigm, researchers present a set of stimuli to remember and then probe one item after a delay, asking participants to report the target by clicking on a circular stimulus report wheel (Fig. 1A). The data are typically analyzed using the circular difference between the true stimulus and reported stimulus, which is then modeled to quantify memory performance^7,8^. Because errors that arise in this task have a “fat tail” — there are more far away errors than you might expect (Fig. 1B) — the dominant models of working memory draw critical distinctions between fundamentally different kinds of memory errors: those caused by limits in how many items are represented vs. how precisely they are represented^7^ or those caused by items encoded with high precision vs. extremely low precision^8^.

**Figure 1.**
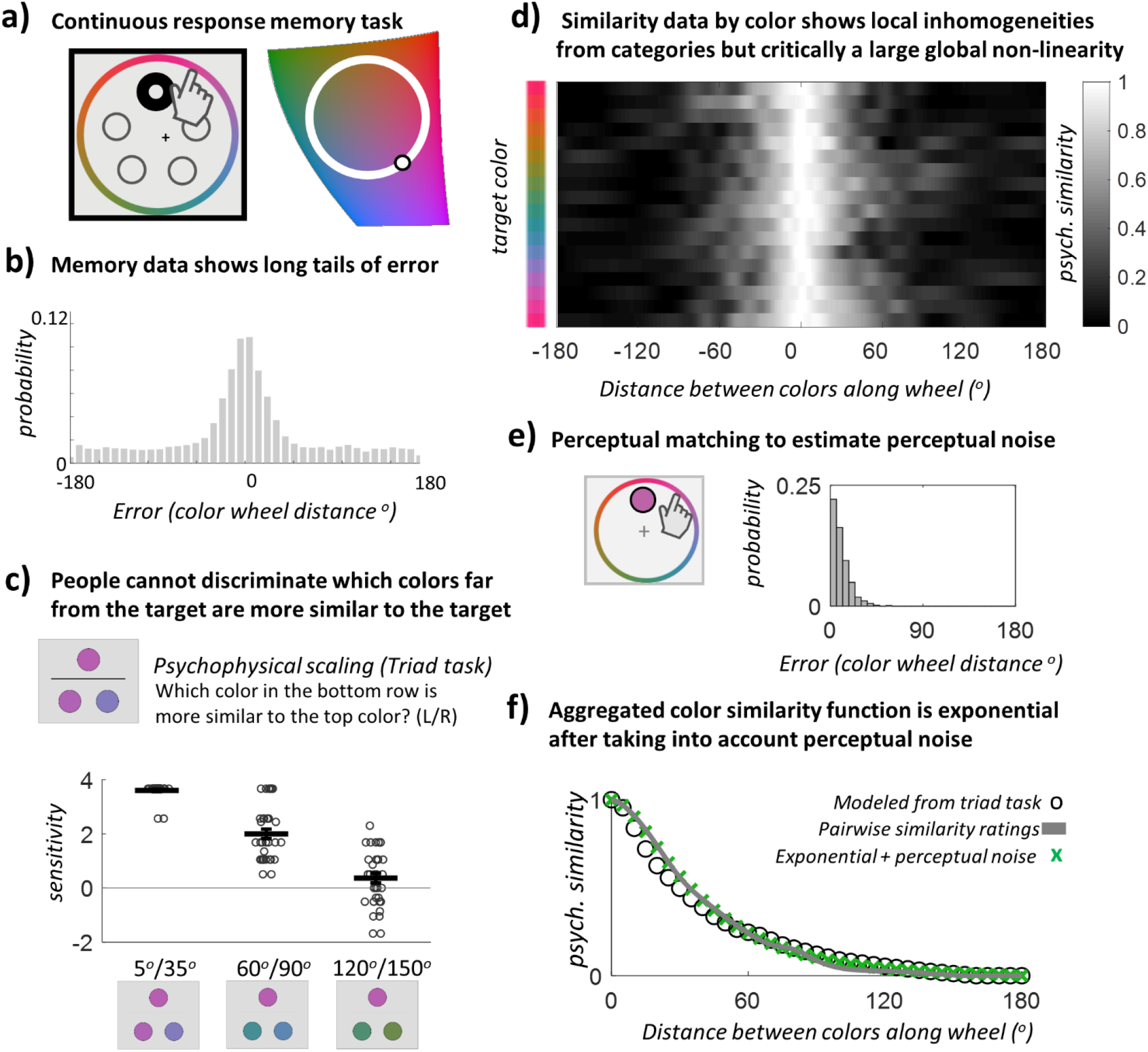
(**A**) A widely used method in working memory is to select a color circle from a slice of color space, show memory items drawn from this circle, and then, at test, probe the contents of a memory item by presenting the entire continuous circle to participants to make a response. Similar response wheels are used for other features, such as face identity. (**B**) A histogram of results generally observed for such tasks, traditionally plotted as a function of distance in degrees of error along the response wheel. There is a ‘long, fat tail’ of errors far from 0 that is often interpreted as evidence for distinct memory states (e.g., guesses or items encoded with very low precision). (**C**) In a triad psychophysical scaling task, N=40 participants had to say which of two colors in the bottom row was more similar to the top (target) color. Despite the difference between the two choice colors always being 30° on the color wheel, sensitivity (d′) dramatically decreased as the choices became more distant from the target, ANOVA F(12,384) = 71.8, p<0.00001, η^2^=0.69. Error bars are within-subject S.E.M. and dots represent individual subjects. See Extended Fig. 1 for the full data. **(D)** We can use the data from another similarity task, a simple pairwise Likert rating of similarity (N=50), to infer the global psychophysical distance of colors at different physical distances along the color wheel. Here we plot this data for sets of target colors, demonstrating previously observed local non-uniformities in color space as the small differences across rows (see Bae et al.^12^).Critically, all of these rows demonstrate a much larger global structure, separate from this local structure: overall similarity falls in an approximately exponential manner. (**E**) Some aspects of this similarity must derive from perceptual discrimination failures (e.g., there are not really 360 independent colors on the color wheel). To estimate this underlying perceptual noise, we use a continuous report task where participants must match a visible color using the same color wheel (N=40) (**F**). We can plot the global psychophysical function -- averaged over all target colors -- using the triad task or the Likert task. Both are very similar and show the same underlying shape. Consistent with previous work, we find this similarity function is exponential once perceptual noise is taken into account (e.g., an exponential convolved with the measured perceptual noise function provides an excellent fit to this data).

Here we present evidence that these small vs. large errors are not distinct kinds of errors, or evidence of multiple psychological constructs being measured (e.g., precision vs. guessing). Instead, we demonstrate that these responses arise fundamentally from a single process. To describe this new conceptualization of memory, we begin with working memory for color as our main case study and then expand the model to encompass working memory for faces (a multi-feature stimulus space) and long-term memory for real-world objects.

The model we propose is a straightforward extension of standard signal detection-based accounts of memory, with the fundamental insight of our framework being the nature of the psychophysical similarity function that explains how familiarity spreads. Consider the simplest case of memory, where you are asked to remember just a single color. When you encode this color — say, red — it will now have significantly enhanced familiarity. Thus, if you are later asked to distinguish the color you saw from a foil color (e.g., red vs. green), the color you saw will likely be more familiar. However, due to noise which corrupts the familiarity signals, this will not always be the case, and on some trials, green might feel more familiar than red.

The critical insight of our model is that when you see red, it does not boost only familiarity associated with red. Instead, a gradient of familiarity will spread to other colors according to a fixed psychophysical similarity function, with considerable activity spreading to similar colors (e.g., pink will also feel familiar), but with much less spreading to dissimilar colors (e.g., yellow, blue and green will experience virtually no boost in familiarity). If asked to hold this color in mind, these initial familiarity signals will be corrupted by noise, and when memory is probed — say, if people are asked to report what color they saw on a color wheel — people will report the color of the response option that currently has maximum familiarity. Although the encoded color is most likely to generate the maximum familiarity signal, competition from other colors (especially from similar colors) ensures that this will not always be the case, and the more noise accumulates, the more likely a very dissimilar color is to be reported. Notably, in this model, memory is not simply a point representation (“I think this item is red”) but instead an entire population of familiarity signals (similar to neural models^9–11^). (We’ve built an interactive demonstration of this model at https://bradylab.ucsd.edu/tcc/ to explain it dynamically.)

According to the model, the way familiarity spreads is a fixed perceptual property, one that can be independently measured using a conventional psychophysical similarity function. Once the nature of the familiarity gradient for a given stimulus space is measured, memory is simply modeled by taking this fixed property of the stimuli and adding noise, with the signal-to-noise ratio (*d*’) being the only memory-based parameter of the model. This model thus uniquely explains the complex shape of error data with only a single free parameter (memory strength, *d*’) and permits parameter-free generalization across different tasks (i.e., without any free parameters, using only measured memory strength and similarity values from different participants). Because this model operates in a signal detection framework, as most models of long-term memory do, it also suggests a unified framework can be used to understand the nature of mnemonic representations and decision-making across working memory and long-term memory.

## Results

### Psychophysical similarity

The most critical component of our proposed model is the psychophysical similarity function that explains how familiarity spreads within a stimulus space (e.g., across the color wheel). While previous work has documented local inhomogeneities in the structure of stimulus spaces^12–14^ we were primarily interested in the global structure of similarity: for a stimulus 10 degrees away on the color wheel from a target color (regardless of what the target color is), how similar is this color to the target on average? Thus, we measured how similarity scales with distance measured in terms of degrees along the color wheel (Methods 1). To do so, we tested how accurately participants could determine which of two test colors was closer in color space to a target color using a triad task^15,16^. This is a perceptual task, but it is analogous to the working memory situation where participants have a target color in mind and are asked to compare other colors to that target. We found that with a fixed 30° distance between two color choices, participants are significantly more accurate at determining which color is closer to the target when the two colors are close to the target in color space compared to when they are far from the target (Fig. 1C, Extended Data Fig. 1; ANOVA *F*(12,384) = 71.8, *p*<0.00001, **η**^2^=0.69). In other words, in a purely perceptual task, participants largely could not tell whether a color 120°or 150° from the target was closer to the target, whereas this ask is trivial if the colors are 5° and 35° from the target. This demonstrates a strong non-linearity in perceptual similarity.

To compute a full psychophysical similarity function, we utilized the just-described triad task with additional distance pairs (Methods 2). We then applied the maximum likelihood difference scaling technique^16^ (MLDS) commonly used for perceptual scaling to estimate how differences between color stimuli are actually perceived. The estimated psychophysical similarity function falls off in a non-linear, exponential-like fashion with respect to distance (Fig. 1F). In color space, it is also well-matched by a smoother measure that requires substantially less data, namely, the pairwise subjective similarity ratings of colors at different distances along the color wheel using a Likert scale (Methods 3; Fig. 1F).

While there are also small local inhomogeneities (Fig 1D), we are primarily interested in the fact that the global structure of similarity space is strongly nonlinear, in agreement with decades of work suggesting psychological similarity is globally exponential (e.g., the universal law of generalization^17,18^), with confusions for very similar colors also caused by perceptual noise^19^ (measured here using a perceptual matching task, Methods 4; Fig 1E, F).

A key implication of these similarity scaling results is that the linear axis of error along the response wheel (e.g., −180 deg. to 180 deg.) previously used to analyze working memory capacity does not capture the psychological representation of the stimuli. This poses a significant challenge to existing memory models, as their parameters are derived assuming linear similarity (i.e. treating the axis of error in degrees as a linear scale). However, this axis is not linear even in a perceptual task: Since participants are essentially incapable of discerning whether an item 120° or 180° from the target in color space is more similar to the target, it is not surprising that they confuse these colors equally often with the target in memory.

### Incorporating psychophysical similarity into a signal detection model

Psychophysical scaling formalizes how similar two stimuli are perceived to be and is the first critical aspect of our proposed model. The next aspect is that signals are corrupted by noise, which we formalize using signal detection theory.

In particular, the model we propose here is fundamentally the same longstanding signal detection model used across decades of research on long-term memory and perception^20–22^, modified to take into account psychophysical similarity. The basis of signal detection theory is that when deciding among each of the colors at test, participants rely upon a noisy, cue-dependent familiarity signal for each color, and the color that generates the maximum familiarity signal is selected (Fig. 2). The stronger the maximum signal is, the higher the confidence in the selected color.

**Figure 2.**
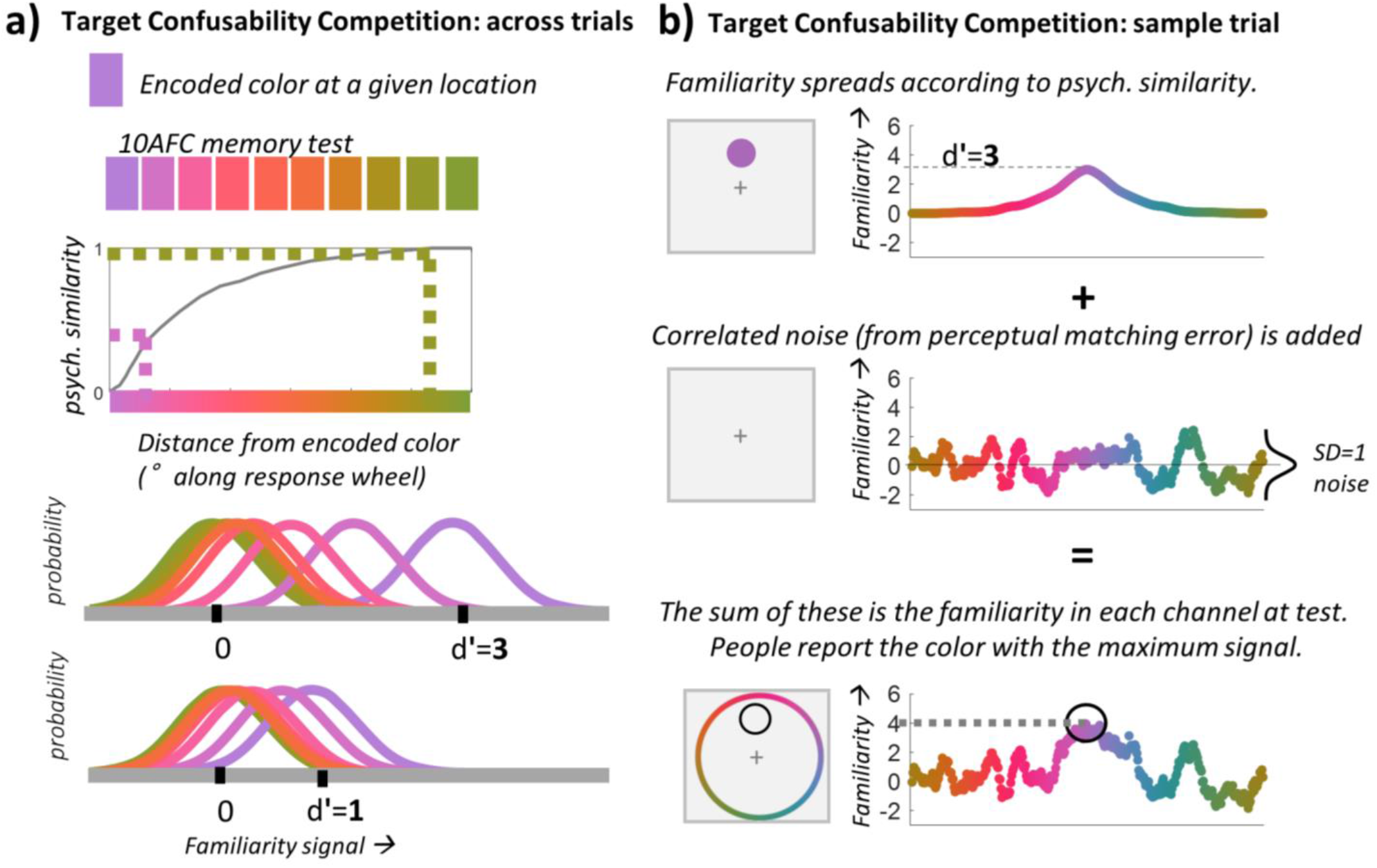
For an interactive version of this figure, see https://bradylab.ucsd.edu/tcc/ (**A**) Our TCC model applied to a hypothetical 10-alternative forced-choice memory test. In standard 2-alternative long-term recognition memory experiments, unseen items vary in their familiarity, which is modeled as a normal distribution. Previously encoded items elicit higher familiarity on average, modeled (in the simplest case) as a normal distribution with a mean of d′, where d′ indicates how many standard deviations of memory strength is added to seen items. When asked what they remember, people pick whichever color elicits higher familiarity on that trial. To generalize to a 10 alternative forced-choice, we thus only need to specify the average familiarity strength of every lure. Usually, all 9 lures are assumed to have a mean of 0 -- with no added familiarity -- when modeling such tasks^21^. However, in a continuous space this is not plausible. Thus, in TCC, we propose that familiarity spreads according to similarity: the mean of each lure’s familiarity distribution is simply its similarity to the target. For example, if the target is purple, other purples will have boosted familiarity as well, and thus people will choose a slightly different purple lure much more often than an entirely unrelated lure such as green. Examples of d′=3 and d′=1 illustrate the idea that when memory for the target color is weaker, more of the lure distributions cluster near the target — and at d′=1, all of the far away colors are in a position to sometimes ‘win the competition’ by having the highest familiarity, but will do so on average equally often, creating a long fat tail. The 10-AFC logic provided here can then simply be adapted to 360-AFC to model continuous report, but with the added knowledge that very similar colors also have correlated noise (measured using the perceptual matching function); i.e., there are not 360 independent colors on the color wheel. (**B**) An alternative way of plotting the same model is to consider a single trial, rather than the distribution of memory strengths across trials. When we encode a purple color, with memory strength d’=3, the familiarity of purple as well as similar colors is increased (according to the measured psychophysical similarity function). Then, we add SD=1 noise to each color channel. The resulting familiarity values, after being corrupted by noise, guide participants’ decisions. In a continuous report task, people simply report the color that generates the maximum familiarity value.

Our model differs from a standard model of the *n*-alternative forced choice only in the usage of the psychophysical similarity measure. In a standard signal detection model of an *n*-alternative forced-choice task, it is generally assumed that exactly one item has been previously seen, so its familiarity is centered on *d*′, whereas the other *n* - 1 items are equally unfamiliar and therefore centered on zero familiarity^21^. However, when memory is tested using a continuous stimulus space, it would be implausible to assume that a color 1° away in color space from the target would have no added familiarity and would have noise that is totally uncorrelated with the target.

Thus, in our model, the mean memory signal for a given color *x* on the color wheel, denoted *d_x_*, is based on that color’s separately measured similarity to the target, i.e., *d_x_* = *d*′ *f*(*x*), where *d*′ is the model’s only free parameter (memory strength) and *f*(*x*) is the empirically determined psychophysical similarity function (i.e. a measurement, done in different participants, of the similarity structure of the color space). The noise added to each color is also correlated between nearby colors according to the empirically measured proportion of how often colors at that distance are confused in a perceptual matching task (Fig 1E), although this is not critical for fitting continuous report error distributions (Extended Data Figure 2).

Because of the nonlinear similarity function, colors in the >~90° physical distance range all cluster near *f*(*x*)≈*f*(*x*)_*min*_ such that *d_x_*≈0 for x~=90° to 180°. Thus, when participants encode a color—say, purple—it increases the average familiarity signal in the purple channel and also in nearby (similar-to-purple) channels while having almost no effect in dissimilar color channels (Fig. 2B). The familiarity signals in each channel are then corrupted by noise, and the resulting reports are based on this noisy signal. In the case of continuous report, people theoretically report the color with maximum familiarity.

Importantly, this Target Confusability Competition (TCC) model can explain all the key features of visual working memory. In particular, it accurately characterizes memory performance across a variety of domains, including different set sizes, encoding times and delays (Fig. 3; Supplementary Figure 1). Previous cognitive models of visual working memory allow for many ways in which memory for an individual item can vary (e.g., guess rate, precision, variation in precision^7–8,23^). By contrast, TCC holds that these experimental manipulations affect only a single fundamental underlying parameter (the memory strength parameter, *d*′), and that the complex changes in the shape of the error distribution arise not from multiple parameters, but simply from the similarity function combined with the non-linearity inherent in selecting only your strongest familiarity value for report. Thus, the fact that manipulations of set size, delay and encoding time — 22 different manipulations in total — result in distributions that can be accurately characterized with only a single varying parameter is strong evidence in favor of TCC, as is the fact that it describes the data extremely well despite being markedly simpler than alternative theories. It is markedly simpler because it proposes a unified generative process for all responses instead of requiring different states to generate different subsets of responses (as in the encoding variability or lack of represented items proposed by previous models^7–8,23^), and because it replaces free parameters (like precision) with independently measured values (like similarity, which is independently measured and fixed for all participants and conditions; Extended Figure 4).

**Figure 3.**
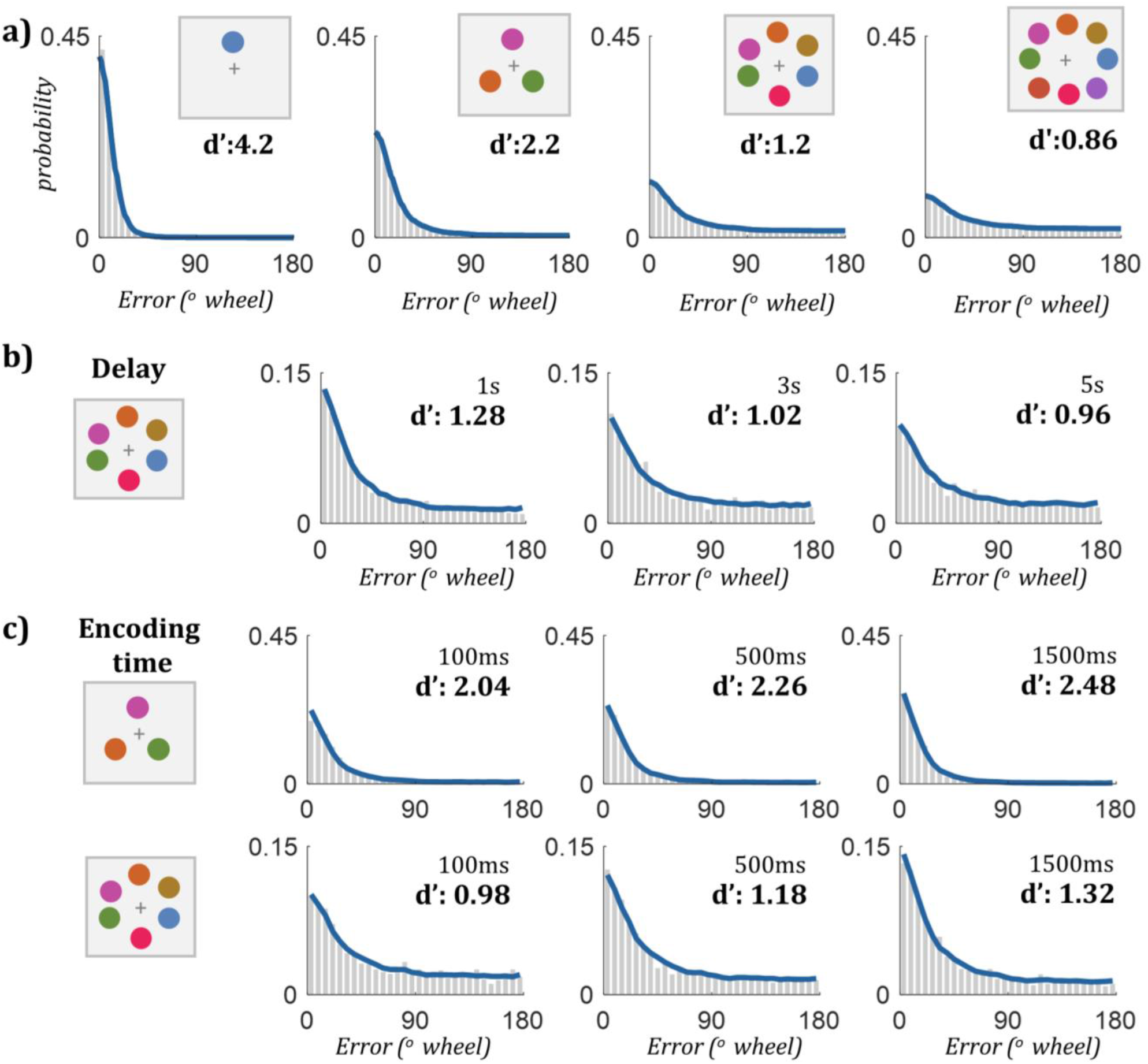
(**A**) TCC fits to group data at set size 1, 3, 6, and 8 (N=20). Despite no concept of unrepresented items or guessing or poorly encoded items, and adopting for the sake of simplicity the assumption that all items are encoded equally (i.e., with the same d’) -- TCC fits even the high set size data accurately because of the noisy nature of the signal detection process combined with the non-linear psychophysical similarity function. (**B**) TCC fits to N=20 group data with varying delay (only set size 6 shown; remainder of data in Supplementary Figure 1). (**C**) TCC fits to N=20 group data across different encoding times (only two set sizes shown; see Supplementary Figure 1). Across several key manipulations of visual working memory (set size, delay, and encoding time), which drastically alter the response distributions collected, TCC accurately captures (with only a single free parameter d′) the response distribution typically attributed to multiple parameters / psychological states by existing frameworks and models of working memory. Only a subset of the delay and encoding time fits are plotted here, but all fits are accurate, as demonstrated by the Pearson correlation between the binned data and model fits as a function of set size (Supplementary Table 1). Note that d’ of the fit to the group data, as plotted, is not the same as the average of individual subject d’s, as used in the model comparisons.

The measured non-linear similarity function is critical to the ability of TCC to fit the data. While reporting the color that is maximally familiar does, on it own, introduce a non-linearity that favors the strongest signals, this alone is not sufficient to explain the data (Extended Data Figure 3). Instead, the explanatory value of TCC comes from the combination of the non-linear similarity function and signal detection theory.

While the main evidence in favor of TCC is its ability to parsimoniously characterize the effects of qualitatively different experimental manipulations (Fig. 3, Supplementary Table 1) and to make precise predictions across tasks and stimuli (see below), we also compared the fit provided by TCC to the fit provided by mixture models of visual working memory, including the standard two-parameter mixture model that interprets performance as arising from distinct concepts of ‘capacity’ and ‘precision’^7^ and a three-parameter version of the mixture model that allows for variable precision^23^. Despite being simpler and having fewer parameters, TCC was just as good at predicting held-out data in a cross-validation test and was reliably preferred in every subject across set sizes when using metrics preferring simpler models (Supplementary Table 2). This was true even though TCC fits are based on aggregated similarity functions from a different group of participants, suggesting the global structure of the psychophysical similarity function is largely a fixed aspect of a given stimulus space. Taking into account color-specific similarity functions (e.g., Fig 1D) or individual differences in similarity scaling should further improve the fit of the model (Extended Data Fig. 5), and would be necessary for comparing the model to others that do take into account such information, but here we focus on the general case of treating all colors and participants as sharing a similarity function.

While memory strength varies according to a variety of different factors (Fig. 3), many researchers have been particularly interested in the influence of set size. TCC shows that at a given encoding time and delay, *d*′—theoretically an interval-scale measure of memory strength^21,24^—decreases according to a power law as set size changes (Extended Data Figure 6), broadly consistent with fixed resource theories of memory^24,25^. Critically, memory strength decreases most at low set sizes (e.g., 1 to 3), suggesting limits of working memory may be best studied across lower set sizes, contrary to the majority of the field which seeks to pressure “capacity” via high set sizes to understand the nature of working memory.

### TCC accurately predicts connections between working memory paradigms that mixture models claim are impossible

Ultimately, evaluating theories based on model comparisons of fit —when all models fit the data well, as here —is not as useful as investigating what they accurately predict^26^. TCC makes a precise and unique prediction that since all responses are generated from the same underlying process, measuring *d*′ in any way that avoids floor and ceiling performance—even using only two maximally dissimilar 180-degree away colors in a 2AFC task —is sufficient to accurately predict (with no free parameters) memory performance involving more similar colors and/or more response options (including continuous report). This is in direct contrast to the inability of mixture models and variable precision models to make such predictions. Such models claim memory varies in multiple fundamentally distinct ways (i.e., precision and guessing can both vary, or the distribution of precisions can vary), and clearly, a single measure of accuracy cannot possibly measure more than one fundamental distinct property of memory.

Specifically, such existing models insist that such predictions should not be possible because they claim that heterogeneity between items is crucial to explaining large vs. small errors. That is, existing models claim that fundamentally distinct items and memory states explain close-to-target responses on the color wheel (e.g., “precision errors for remembered items” or “high precision items”) vs. responses far away from the target (e.g., “guesses” or “low precision items”). Thus, existing models inherently assume that a singular measure of how well participants can discriminate 180°changes (e.g., was it red or green?), which measures only information about items that cause large errors, cannot, even in principle, measure the properties of the items that cause small errors. By contrast, TCC says all responses to more similar colors are directly predictable using the fixed similarity function, and that memory varies in only one way (memory strength), and thus such a 2-AFC task is sufficient to measure memory performance.

In two experiments, we tested TCC’s prediction that a single measured *d*′ is sufficient to characterize memory performance across a variety of tasks that are currently thought to tap different memory processes. In both experiments we had participants perform a memory task involving a 2-AFC test with maximally dissimilar colors (two options: 0° away from the target color vs. 180° away from the target color). We used the data from this 2-AFC task to compute *d*′ in the standard way (denoted *d*′_180°_) and then used TCC —with this exact *d*′ —to compute parameter-free predictions for a variety of other conditions. We intermixed all the conditions – including conditions that require participants to remember the precise color they saw -- so that participants could not rely on a categorical memory strategy in the maximally distinct 2-AFC task.

In one experiment involving a 2AFC task (Fig. 4), we used TCC with fixed *d*′_180°_ to predict how well participants could discriminate the target from more similar foils (e.g., to predict *d*′_12°_ from a 2-AFC task involving the color they saw vs. a color only 12° away). With no free parameters, memory performance was accurately predicted over the entire range of intermediate foil similarities (Fig 4C). TCC accomplished this with no free parameters because it specifies how the perceptual similarity of the two colors on a 2-AFC task (measured in a separate psychophysical procedure) should impact memory performance (see also Kahana & Sekuler^27^; Nosofsky^19^). By contrast, mixture models, based on the distinct concepts of guessing and precision, anticipate no particular relationship between performance on a 2-AFC task involving maximally dissimilar foils and performance on a 2-AFC task involving more similar foils. 2-parameter mixture models can use 180° 2-AFC performance only to measure ‘guess rate,’ leaving precision unspecified. Thus, with only 180° 2-AFC performance in hand, these models are able to predict a wide range of possible outcomes on 2-AFC tasks with more similar foils, depending on the unknown factor of ‘memory precision’ (Supplementary Figure 2). Note that precision, unlike similarity, is thought to be changed by memory strength and differ across subjects, and thus precision measures are not constrained by fixed perceptual similarity data that TCC can utilize so effectively. Because the mixture model predictions are largely unconstrained, TCC is strongly preferred to mixture models by a Bayes factor model comparison (group Bayes factor preference for TCC > 200:1, individual subjs: t(54)=11.19, p<0.001, d_z_=1.51, CI=(2.9:1, 4.2:1)).

**Figure 4.**
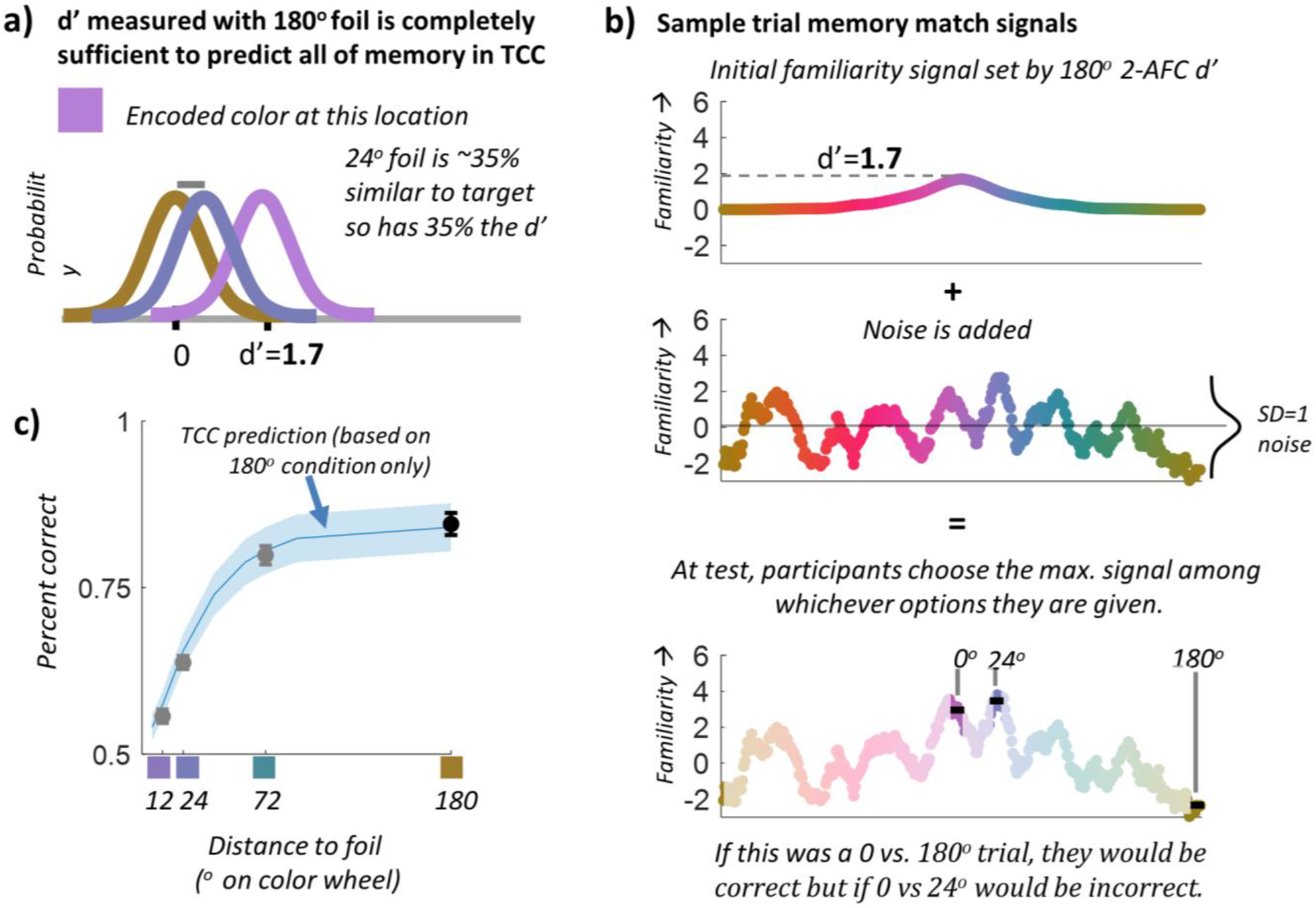
(**A**) Since TCC states that visual working memory performance is determined by simply d′ (memory signal strength) once perceptual similarity is known for a given feature space, it makes novel predictions no other theory of working memory can make. In particular, it predicts that d′ measured with a 180 degree, maximally dissimilar foil (i.e., d′_180°_) should be completely sufficient to predict all of memory performance, unlike models where errors to maximally dissimilar foils arise from different processes than errors to similar foils, e.g., where errors to maximally dissimilar foils solely from ‘guessing’ (in some models) or from extremely poorly encoded items (in other models). For example, after measuring d′_180°_, TCC predicts that since a 24 degree foil is ~35% similar to the target, discriminability on a 2-AFC task in which the foil is 24 degrees away from the target should be ~35% of d′_180°_. (Although note that correlated noise makes this more complex for very similar foils) (**B**) On a single trial, this prediction can be visualized in a straightforward way: If we know the target was encoded with d′_180°_ = 1.7, then TCC makes a strong prediction about how this familiarity spreads to other colors and how it is corrupted by noise. In continuous report, the decision rule is to report the maximum of the resulting color channel familiarity responses; in 2AFC, the decision rule —based on the exact same underlying color channel responses —is to choose the highest familiarity signal of your response options. Thus, in this example trial, the participant in a 2AFC task would choose the 0°-target over a 180°-foil, but would choose a 24°-foil over the 0°-target. Because TCC specifies this entire generative process, it makes precise predictions about how often people will make errors to different distance foils. (**C**) Here we plot the predicted percent correct of different distances of colors from the target (blue), a prediction based only on performance from the 180 degree condition (black) with no free parameters. When comparing subject’s performance at different foil distances (gray, N=60) we demonstrate TCC accurately predicts performance across different foil distances.

In a second experiment we went further, showing that TCC — again using only measured *d*′_180°_ from a 2AFC task and separately measured perceptual similarity between the response-option colors in different participants — can accurately predict performance when there are more than two response options, up to and including continuous report, again with no free parameters (Figure 5). In this experiment, we once again found a strong preference for TCC’s prediction over the mixture model models in generalizing from 2-AFC to continuous report, which is the only condition the mixture model can be fit to (group BIC preference for TCC > 650:1, individual subjects: t(51)=7.64, p<0.001, d_z_=1.06, CI=(9.5:1, 16.2:1)). We also found that 2-AFC *d*′ measured in the standard way (i.e., *d*′_180°_) maps directly to TCC’s *d*′, which explains the full continuous report distribution (Fig. 5B). The lopsided Bayes factors arise because TCC precisely predicts the outcomes (outcomes that, when tested, are empirically observed), whereas competing models necessarily claim that the 2-AFC data are insufficient to completely measure memory since they do not measure the ‘precision’ of memory.

**Figure 5.**
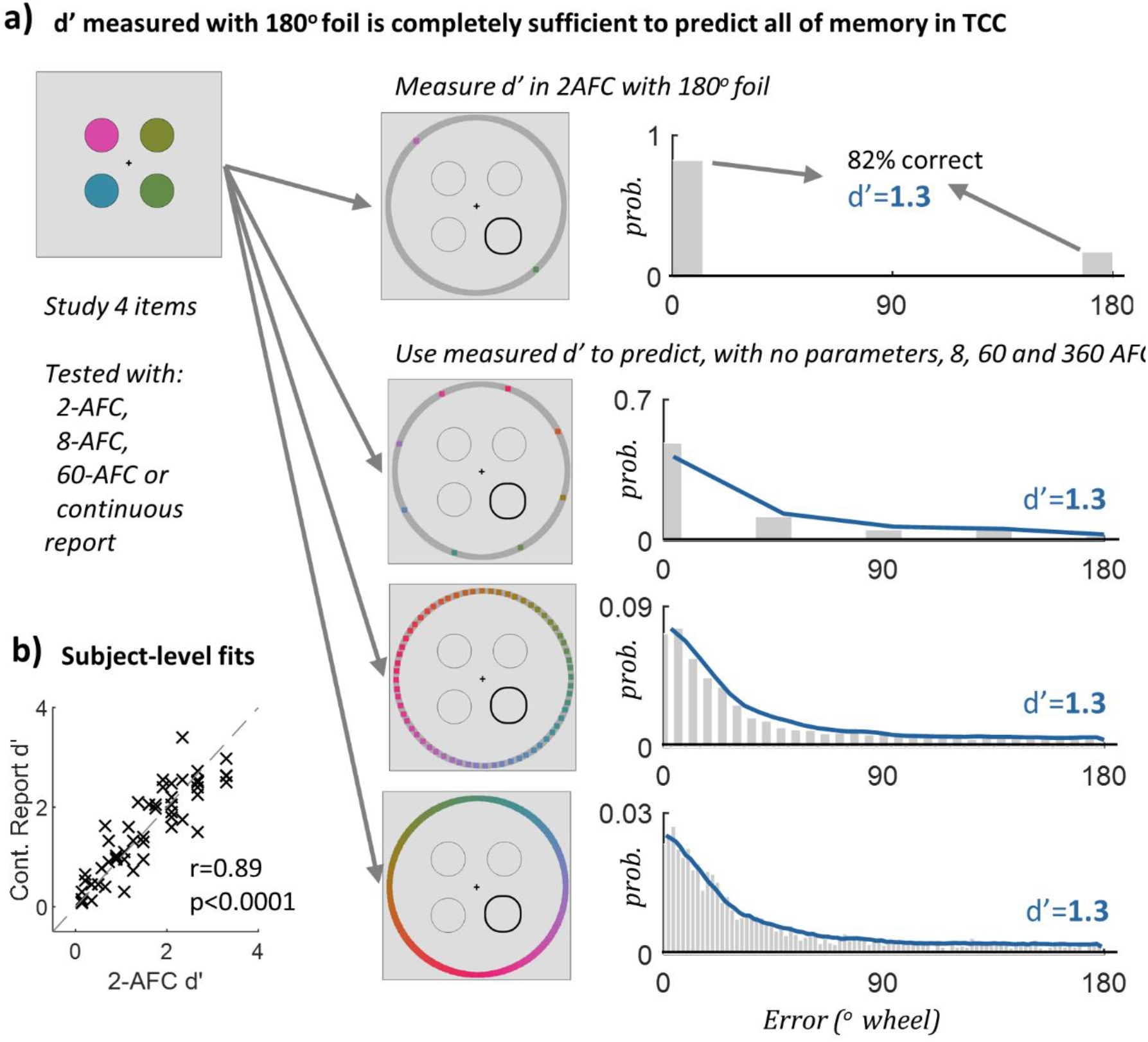
(**A**) According to TCC, the d′ in a 2AFC task is fundamentally the same d′ in continuous report tasks (or any other AFC task). Thus, unlike other models, TCC makes a strong prediction that d′ as measured with a 180 degree foil (d′_180°_) is completely sufficient to predict all of memory across any number of options presented at test, including completely sufficient to predict the entire distribution of errors in continuous report (since ultimately this distribution does not arise from distinct psychological states, but simply from combining the fixed similarity structure of the stimulus space with memory strength). To test this prediction, N=60 participants encoded items into memory and were then tested using 2-AFC, 8-AFC, 60-AFC or continuous report (360-AFC). During 2AFC trials, the foil was always 180 degrees away, which we used to calculate d′_180°_. We then used TCC, with this measured d′ but with no free parameters, to accurately predict 8-, 60-, and 360-AFC performance. The accuracy of these predictions provides further evidence there is no need for forgotten or low-precision items to account for the tail of continuous report distributions. Instead, for a given stimulus space, the continuous report distribution is modulated by memory strength but is otherwise always the same shape, determined by the shape of the similarity function for that stimulus space. (**B**) We can also independently estimate d′ from the continuous report data and from the 2-AFC data. We find a strong subject-level correspondence between TCC’s continuous-report estimate of d′ and d′ estimated from the 2-AFC task in the traditional way (i.e., d′_180°_), Pearson r=0.89, p<0.001, CI=(0.81,0.93), in line with what is expected simply from the noise ceiling of these measurements. Each point is a subject mean.

Thus, with TCC, measuring only how well participants can distinguish between far apart test items (0° vs.180°) using a 2-AFC task is sufficient to predict the distribution of responses from a continuous report task and to predict 2-AFC performance for distinguishing targets and foils of varying similarity (so long as the 2-AFC task is not at ceiling or floor). Together, these experiments provide compelling evidence against previous models of visual working memory where the tails of the continuous-report distribution (the only aspect of performance that is theoretically measured with 180° foils in 2-AFC) are fundamentally distinct from the center of the distribution.

In other words, in the competing models, responses in the tails of the distribution result from ‘guesses’ or ‘low precision’, whereas the central responses result from high precision memories. If these models were correct, it should not be possible for TCC to make such accurate predictions across tasks using a single *d*′ and no free parameters. The fact that TCC can make such accurate predictions allows the reintegration of a huge literature on change detection with very distinct foils, with important theoretical and clinical implications^28^, as it shows that measuring *d* with maximally distinct foils is sufficient to understand memory response distributions —there is no separate “precision” that is not being measured in such tasks.

### Generalization across different stimulus spaces

So far we have focused largely on color space, which is the dominant way visual working memory is studied^7^. However, TCC is not limited to color and can be applied to any stimulus space. To demonstrate its generality, we applied TCC to the case of face identity, since it is a complex stimulus space that contains multiple low- and high-level features. Using a previously created face-identity continuous report procedure^29^, we collected memory data for set size 1 and 3. We also measured the psychophysical similarity function and measured the accuracy of perceptual matching on this face space (Fig. 6). Again, we found the TCC fit observed memory data extremely well across both set sizes 1 and 3 (see Fig. 6) and fit reliably better than existing mixture models (Supplementary Table 3).

**Figure 6.**
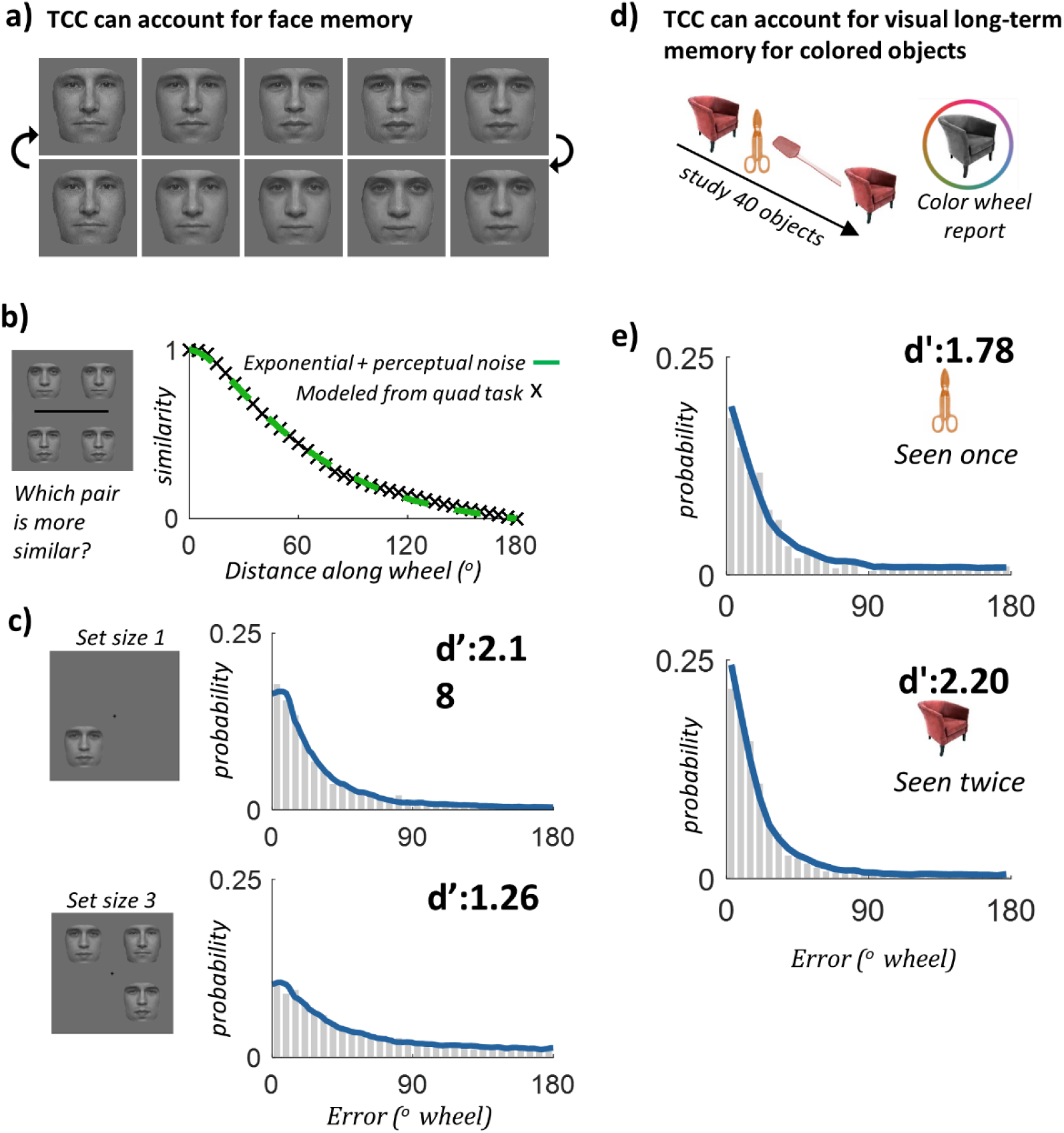
(**A**) Examples from a previously used continuous face space^29^. (adapted from^29^). **(B)** Using a ‘quad’ similarity task to reduce relational encoding, and the same MLDS method and perceptual matching task as with color, we collected a psychophysical distance function for face identity, N=102. (**C**) TCC fits to working memory data (N=50) using face identity at set size 1 and 3 (r=0.997, p<0.001, CI=(0.993, 0.998); r=0.985, p<0.001, CI=(0.971, 0.992). TCC accurately captures face identity data, demonstrating its generalizability across diverse stimulus spaces. **(D)** To show generalization to other memory systems, we fit data on a visual long-term memory continuous report task with colors^30^. N=30 participants performed blocks of memorizing 40 items, and then after a delay, reported the colors of the items using a color wheel. Some items were seen only once, and some repeated twice in the same color within a block. **(E)** TCC fits to visual long-term memory data for items seen only once and for items repeated twice (r=0.978, p<0.001, CI=(0.958, 0.988); r=0.991, p<0.001, CI=(0.983, 0.995)). TCC accurately captures visual long-term memory data, suggesting the psychological similarity function is a constraint on both working and long-term memory systems. Note that long-term memory performance in this task likely depends on a two-part decision —item memory and source memory (e.g., the object itself, and then its color). This two-part decision is related to the processes of recollection and familiarity and likely introduces heterogeneity in memory strength into the color memory reports. Here, where item memory was consistently strong and color memory was the main factor, this does not affect the fits of TCC, but in other data where heterogeneity in strength of item memory is greater, variability in d’ between items would likely need to be accounted for.

Thus, TCC accounts for data across multiple stimulus spaces. As long as the perceptual similarity space of the stimuli is accurately measured using psychophysical scaling (see Supplementary Discussion), TCC’s straightforward signal detection account, with only a single *d*′ parameter, accurately captures the data.

### Generalization across different memory systems

To demonstrate TCC’s applicability to multiple memory systems, not just visual working memory, we fit data from a visual long-term memory continuous report task with colors. Unlike the previous datasets, this data had been previously reported in the literature^30^. Participants performed blocks where they sequentially saw 40 real-world objects’ that were randomly colored, and then after a delay, reported the color of the object using a color wheel (as in Brady et al.^31^). Some items were seen only once, and some repeated twice in the same color within a block (Fig. 6D). Again, we found that TCC fit the observed memory data extremely well across both the unrepeated and repeated items (Fig. 6E). Thus, unlike working memory modeling frameworks which propose system-specific mechanisms (e.g., population coding combined with divisive normalization^10^), TCC naturally fits data from both visual working memory and long-term memory with the same underlying similarity function and signal detection process applicable across both memory systems.

### Implications of TCC: no objective guessing

One particularly important implication of TCC’s fit to the data with just a single parameter is that it implies there is little-to-no objective “guessing” in working memory. This provides evidence against a fixed capacity limit where participants only remember ~3-4 items^1,2^ and is consistent with more continuous conceptions of working memory^4^. In particular, while colors far from the target in color space sometimes ‘win the competition’ (e.g., have maximal familiarity after noise is added), this is not because the target was fundamentally unrepresented or varied hugely in encoded memory strength trial to trial. In a stochastic competition, the strongest representation does not always win. Moreover, the target will be more likely to lose the competition the weaker its representation is. Critically, in TCC, at least as proposed so far, the target is always represented —that is, people’s familiarity signals are never unaffected by what they just saw 1 second ago (as in *d*′=0).

While these conclusions follow from the excellent fits of the straightforward 1-parameter TCC model to a wide variety of data (data widely thought to provide prima facie evidence for the existence of unrepresented items) and from the generalization of maximally-dissimilar 2-AFC performance to other conditions, to evaluate this claim further, we assessed a 2-parameter hybrid model based on TCC but mixed with objective ‘guessing’. This hybrid model assumes only a subset of items are represented and that the remainder have *d*′=0. Focusing on the highest set sizes (6 and 8), we found such a model was dispreferred in model comparisons in 100% of subjects compared to TCC (BIC, set size 6: t(19)=-41.99, p<0.001, d_z_=9.39, CI=(6.2:1, 6.9:1); t(19)=-16.09, p<0.001, d_z_=3.60, CI=(5.3:1, 6.9:1)), and BIC was well calibrated for these model comparisons (Supplementary Figure 3). Furthermore, while this hybrid model accurately recovered its own parameters from simulated hybrid data, showing it detects objective ‘guessing’ if it is present (Fig. 7C), when fit it to empirical data it estimates ‘guessing’ rates near 0 in every set size in group data (Fig. 7B), and a guess rate <5% in the majority of individual subjects at every set size. Thus, although some items may occasionally have a *d*′ of 0 (perhaps because they were completely unattended during encoding), it appears to happen too infrequently to appreciably affect the fit, and it happens far less often than required for ‘slot’ models of working memory that suppose 4-5 of the 8 items are always entirely unrepresented^2^. The simulation results demonstrate it is possible to detect “random guesses” if present in the data, but TCC finds no evidence for such objective ‘guessing’ in real data. Critically, however, like any standard signal detection model, TCC naturally accounts for the subjective feeling of guessing/low confidence^21^ that arises when memories tend to be weak, like at high set sizes (Extended Data Fig. 7 and 8).

**Figure 7.**
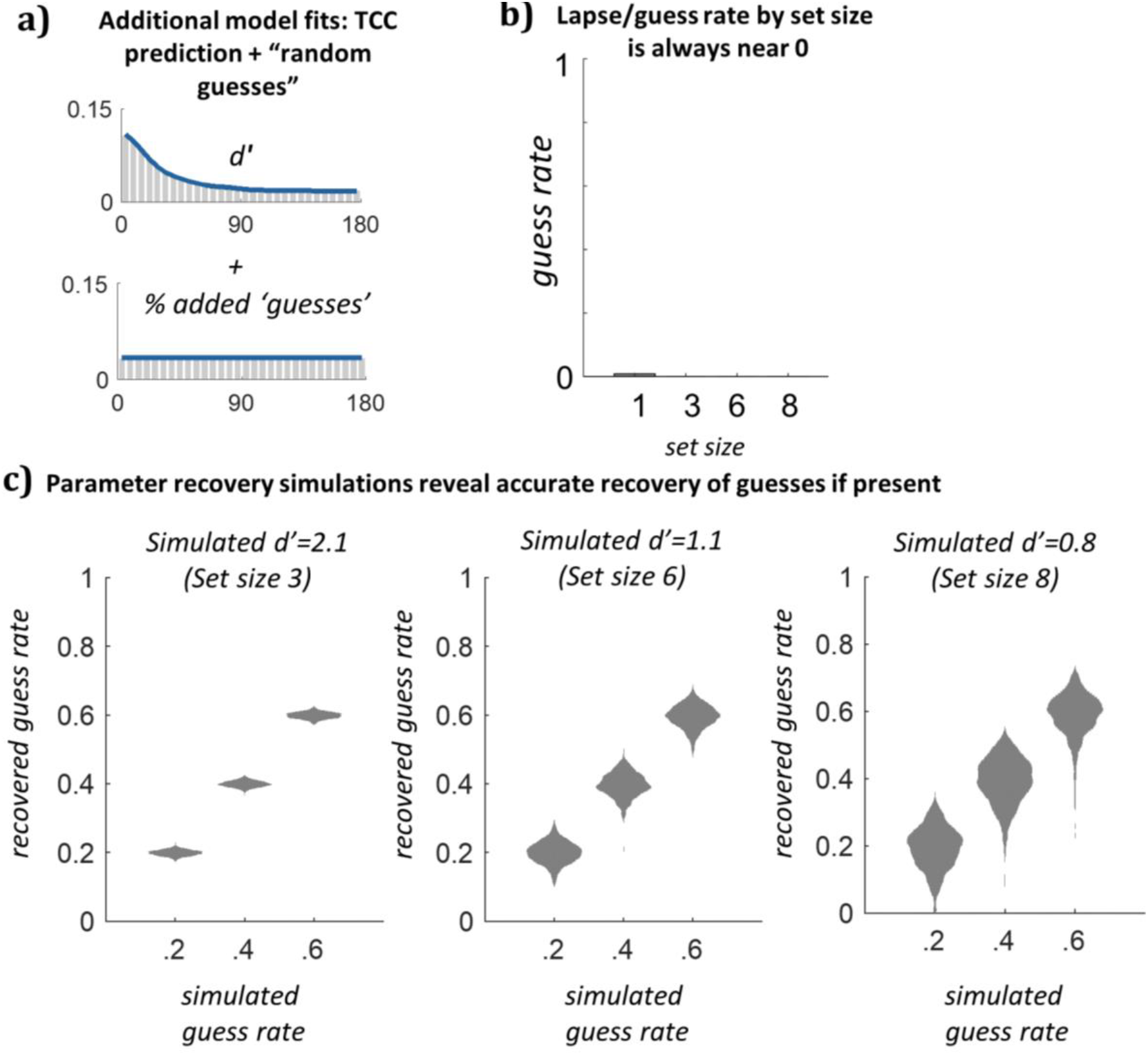
(**A**) To validate whether TCC could detect objective guessing (i.e. a separate psychological state with no information) if present in the data, we considered a mixture of responses from TCC plus objective guessing, creating a mixture model of TCC and a uniform distribution. **(B**) Although model comparison strongly preferred TCC with no guessing, we nevertheless fit a hybrid TCC+guessing model (2 parameters) to real participant data, and found that the guessing parameter in real data is estimated at ~0 across all set sizes. (**C**) However, when fitting the hybrid TCC+guessing model to simulated data, we observed accurate recovery of guessing if present in the data -- even 20% ‘guesses’ added to set size 8 d′ levels is accurately recovered and never estimated as 0 (data are violin plots, showing entire distribution of recovered parameters). Furthermore, model comparison metrics —even those, like BIC, designed to prefer simpler models— prefer the hybrid model with the guessing parameter in every simulation with guessing added (all BIC>30:1 in favor of hybrid model). This provides strong evidence there is little objective ‘guessing’ in visual working memory data, and that our modeling with TCC would be able to detect any significant number of added no-information responses if they were present.

### Implications of TCC: mixture models are not measuring distinct psychological states

The dominant quantitative model of visual memory is the mixture model, which claims to measure two distinct psychological concepts from continuous report error data: how precisely people remember items they have in mind (e.g., “precision”; “variability in precision”) and how often people have an item in mind (“likelihood of retrieval”, or its opposite, “guess rate”). The fundamental claim that there are two distinct ways memory can fail —loss of precision or loss of discrete items —permeates a huge variety of literature in working memory, attention^32^, iconic memory^33^ and long-term memory^34^. TCC makes a counterclaim: the fact that manipulations of set size, delay and encoding time that hold the stimulus space constant (e.g., use a particular color wheel) can be fit by varying a single memory strength parameter; and that measuring how well people can distinguish only maximally distinct comparisons (like red vs. green) is sufficient to characterize memory appears to falsify the idea that memory changes in two or three psychologically distinct ways (e.g., precision vs. guess rate). Another way to test this is to fit the mixture model —which purports to measure two distinct parameters —to data from a single stimulus space (e.g., from a single color wheel) and ask whether the state-trace plot shows evidence of a single way memory changes or multiple ways^35^.

Figure 8 shows this plot for all data from the current paper (e.g., the 22 conditions shown above, plus the other experiments) and from all the conditions in Miner et al.^30^, which provided the long-term memory data fit above. As can be clearly seen in this plot, the two parameters always change together: while not linear in their relationship, they are nearly perfectly related —and their relationship is well predicted by the zero-free-parameter prediction of TCC (e.g., TCC’s prediction across a range of *d*′ values). The non-linear relationship accounts for most cases where people have found evidence to “dissociate” the two parameters (see Supplementary Discussion). This is further evidence that TCC’s single parameter conception of performance is correct and that mixture models are not measuring distinct psychological constructs (see also Supplementary Figure 4 and 5 and Supplementary Table 4, which use data from the literature, although not holding the stimulus space constant as here).

**Figure 8.**
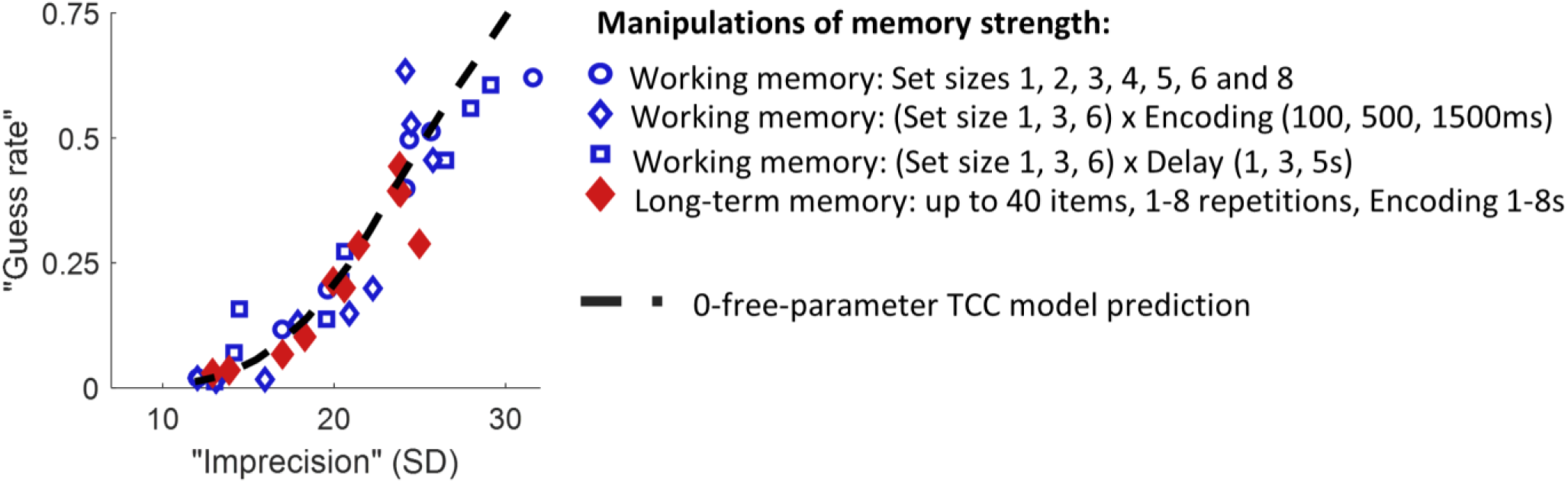
The currently dominant conception of memory arises from mixture models which claim that memory varies in at least two psychologically distinct ways: the precision of memory and the number of represented items (modeled as “guess rate”). TCC makes a strong counter prediction: that if the stimulus space, and thus psychophysical similarity function, is held constant, memory report distributions vary in only one way, in memory strength. Thus, TCC claims that the particular manipulation (encoding, set size, delay) used to change memory strength should not selectively change one mixture model parameter or another (e.g., encoding changing precision; high set sizes affecting only guess rate, etc), but that both should always change together. To visualize this, we show a state-trace plot of mixture model parameters across a wide range of manipulations of working memory (from the current paper) and long-term memory (from Miner et al.^30^), with one point per condition. We find that despite the huge number of different ways we vary memory strength, all the points lie on a single line, consistent with only a single parameter being varied; and that this line is extremely well predicted by the 0-free-parameter prediction of TCC. TCC can only predict an extremely small part of the possible space the mixture model can predict, and only a very particular relationship between the two mixture model parameters, and the data from all of these conditions land on this line. This provides strong evidence against mixture models measuring two distinct parameters and in favor of the TCC conception of memory.

## Discussion

Most previous theories and models of visual working memory have not considered the relationship between stimuli and the psychological similarity of those stimuli. In the absence of psychophysical scaling and without regard for its theoretical implications, the use of these models has led to what we show are illusory ‘independent’ estimates of ‘guessing/capacity’ and ‘precision’ and to arguments for limited capacity characterized by so-called “discrete failures” of working memory, attention^32^, iconic memory^33^ and long-term memory^34^. Indeed, claims about selective deficits in clinical populations^36–38^, and even about the nature of consciousness^32^ have been made based on dissociations between model-based estimates of ‘precision’ and ‘guessing’. Here, we have shown these apparent dissociations are an illusion of modeling the data without taking into account the non-linear way the familiarity spreads in stimulus space. When this fixed perceptual similarity structure is taken into account, TCC provides a unifying theory of visual memory strength, one that is capable of bridging distinct tasks and stimulus conditions that would not be possible using previous models and that undermines the interpretation of apparent ‘discrete’ failures of attention and memory^32–34,36–38^.

While TCC rejects the idea that the distribution of responses collected from continuous report is explained primarily by remembered and not-remembered items (or items that are encoded with extremely different precisions^8^), this does not mean some variability between items is not present in working memory tasks. Psychophysical scaling can naturally account for many stimulus-specific variability effects (e.g., some colors being more distinct than others, Extended Data Fig. 5) by using separate similarity functions for each target color. Furthermore, in light of the signal detection framework of TCC, much of the existing evidence for ‘variable precision’ does not actually provide direct evidence of variability in the *d*′ parameter of the TCC model. Many aspects of variability between items arise in TCC naturally from the independent noise added to different items that is at the heart of signal detection theory, such as the effect of varying confidence on continuous report data or allowing participants to choose their best item for report (Extended Data Fig. 7 and 8). Thus, it remains an open question to what extent *d*′ varies between items and trials. In TCC, if such variation needs to be accounted for, this would be done by moving to an unequal variance signal detection model, whereas the current modeling has used a purely equal variance model. Critically, however, we show that mixing in items that are unrepresented (*d*′=0) is inconsistent with the data. Thus, any variability in *d*′ that does exist across items likely does not include an appreciable role for items with *d*′ = 0.

Many models of working memory focus almost exclusively on how memory strength changes with set size, taking this as the central factor in how much understanding of working memory they have achieved. We take a fundamentally different view, seeing our measure of memory strength (*d*’) as a measure of signal-to-noise that is likely modulated by many factors, and which has a shared structure not only in working memory, where set size matters so much, but also in long-term memory, which appears to follow fundamentally the same rules of memory confusability and a similar decision process (Fig. 8; Miner et al.^30^). Notably, we find that while set size modulates memory strength in the current work, there are many other factors that affect memory strength nearly as much. For example, increased delay decreases *d*′ (more noise accumulates even with the same ‘signal’), and increased encoding time improves *d*′ (more signal relative to the same noise). Similarly, in some situations other factors like location noise (e.g., “swaps”; Bays et al.^33^) and ensemble coding^40,41^ seem to play a major role in memory errors. Thus, while we find an approximately power law-like relationship between set sized and d’ (Supplementary Figure 4), we are hesitant to assume that there will be a fixed ‘law’ for how set size relates to memory errors and note that previous work that claims to find such rules^7,8^ has almost never examined whether those rules hold when manipulating other factors that will also independently impact memory strength, like encoding time and delay.

In addition, in the current work, we present a straightforward version of the TCC model that does not account for all possible factors. For example, it is possible to make different predictions for different target colors, taking into account category effects (e.g., Extended Data Fig. 5). In addition, while in the current data we see almost no ‘swaps’ or location-based confusions (because we use long encoding times and placeholders), it is of course possible to implement a ‘swap’ parameter in TCC (as in Williams, Brady & Stoermer^42^) or explicitly model the psychophysical similarity structure of location and therefore make parameter-free location confusion predictions. Similarly, hierarchical models of ensemble coding and grouping, or other forms of integration across items could potentially be implemented using TCC as the basis of memory responses. If there is significant integration across items or across time in a particular paradigm, more complex models like these would be needed because TCC’s item-based prediction about error distributions would no longer be a valid assumption.

While TCC is a theory about the fundamental nature of the underlying memory signal in visual working and long-term memory tasks, and about how this signal is used to make decisions, there are many potential cognitive and neural explanations (shared or independent across systems) that may instantiate the model. Indeed, in long-term memory, signal detection models have often been conceptualized in relation to neural measures, including both neuroimaging^43^ and single-unit recording^44^.

The central feature of TCC is the psychophysical similarity measurement, which provides the basis for the straightforward signal detection model. This similarity function is naturally understood using models of efficient coding^18^ or population coding^10^. For example, the idea that far away items in feature space are all approximately equally similar arises naturally from population codes —if individual neurons’ tuning functions only represent a small part of color space (e.g., 15° on the color wheel), there would be extremely limited overlap in the population of neurons that code for any two colors even a medium distance apart on the wheel. There would also be correlated noise between nearby colors, as we assume in TCC.

Thus, the current model is in many ways related to existing models of working memory based on population codes^9,10^. Indeed, the similarities between the framework of population coding and the cognitive model proposed here offers significant promise for bridging across levels of understanding in neuroscience, with a population coding implementations of TCC possible^45,46^. However, as compared to existing population-based models^10^, the cognitive basis of the current model —with the measured scaling function following the well-known cognitive laws of similarity^17,19^ —allows us to fit data with an extremely simple 1-parameter model that allows generalization across tasks and draws strong connections to signal detection theory and long-term memory that are not apparent when thinking about population coding alone without this cognitive basis. In addition, framing our model in terms of signal detection theory allows a very general model of the decision process, compared to population coding models where the decision process is based on variability in spikes in a fixed neural population^22^, which are hard to reconcile with data from high-level stimuli like faces, which are likely encoded in many distinct populations, and data from long-term memory, which is not stored ‘online’ in a fixed neural population.

Previous work has shown psychophysical similarity metrics are likely distinct for different stimuli in the same stimulus space (e.g., memory varies across colors^12,13^; Extended Data Fig. 5). The underlying space upon which the exponential similarity function is imposed may be designed to take advantage of efficient coding of environmental regularities^47^, such that the more frequent the stimuli, the more neural resources we devote, giving improved discriminability and predictable memory biases^48^. Taking this into account may allow a simple parameterization of not only the average similarity function, but the particular functions for individual stimuli (as in Fig. 1D). In addition, psychophysical similarity may not be a fixed property but may be dependent on how the current environment affects discriminability^49,50^. For example, memory biases are altered when discriminability is affected by adaptation or contextual effects^48^.

Some previous models of visual working memory have, like TCC, rejected the idea that the ‘fat tails’ in the error distribution (Fig. 1) arise from unrepresented items^8,9^. For example, models like the variable precision model^8^ hold that items vary in the precision with which they are encoded, and this heterogeneity between items is critical to explaining the shape of the error distribution; i.e., extremely poorly represented items, rather than completely unrepresented items, explain the tail of the error distribution. Like TCC, this model holds that there is not in fact a completely uniform, flat tail in the distribution; and assumes that items vary in representational fidelity (i.e., the independent noise for different items in TCC).

However, in other ways, the two models differ substantial. The variable precision models, like other previous memory models, relies on the assumption that the response axis can usefully be thought of as linear. By contrast, we have shown that similarity and memory confusability are deeply non-linear along this axis, in agreement with decades of work suggesting psychological similarity is globally exponential (e.g., the universal law of generalization^17,18^). This results in significant differences between the variable precision model and TCC. In particular, in the variable precision model, the latent distribution of precisions is an unknown that is taken to vary between situations, whereas TCC uses the insight that similarity is non-linear and relatively fixed to greatly simplify the model of the error distribution (allowing, for example, the generalizations from 180° 2-AFC that are not possible in the variable precision model).

Finally, TCC provides a compelling connection between working memory and long-term recognition memory, which is often conceptualized in a signal detection framework. In particular, it can be naturally adapted to explain a number of findings that are in common between the working memory and long-term memory literatures but have been difficult to explain with previous working memory models, like the relationship between confidence and accuracy^52,53^ (Extended Data Fig. 7 and 8) and the ability of participants to respond correctly when given a second chance even if their first response was a ‘guess’ or ‘low precision response’^51^. Thus, despite research on working and long-term memory operating largely independent of one another, TCC provides a unified framework for investigating the distinctions and similarities in memory across both domains by emphasizing that competition and perceptual confusability between items is a major limiting factor across both working memory and long-term memory.

## Methods

All conducted studies were approved by the Institutional Review Board at the University of California, San Diego, and all participants gave informed consent before beginning the experiment. All color experiments used a circle in CIE *L*\**a*\**b*\* color space, centered in the color space at (*L* = 54, *a* = 21.5, *b* = 11.5) with a radius of 49. All sample sizes were decided a priori, and are similar to those in previous publications^7–9,31^. Approximately half of the data comes from experiments run in the lab, with the others were conducted using Amazon Mechanical Turk. Mechanical Turk users form a representative subset of adults in the United States^54^, and data from Turk are known to closely match data from the lab on visual cognition tasks^40,55^, including providing extremely reliable and high-agreement on color report data^41^. Any systematic differences between the lab studies – where we collect most memory data – and the Turk studies – where we collect most similarity data – would decrease the appropriateness of the similarity function for fitting the memory data, hurting the fit of TCC. Data collection and analysis were performed with knowledge of the conditions of the experiments. All statistical tests are two-tailed.

### 1. Fixed distance triad experiment

N=40 participants on Amazon Mechanical Turk judged which of two colors presented was more similar to a target color. The target color was chosen randomly from 360 color values that were evenly distributed along a circle in the CIE *L*\**a*\**b*\* color space, as described above. The pairs of colors were chosen to be 30 degrees apart from one another, with the offset of the closest color to the target being chosen with an offset (in deg) of either 0, 5, 10, 20, 30, 40, 50, 60, 70, 80, 90, 120, 150 (e.g., in the 150 degree offset condition, the two choice colors were 150 and 180 degrees away from the target color; in the 0 deg offset condition, one choice exactly matched the target and the other was 30 deg away).

Participants were asked to make their judgments solely based on intuitive visual similarity and to repeat the word ‘the’ for the duration of the trial to minimize the use of verbal strategies. Each participant completed 130 trials, including 10 repeats of each of the 13 offset conditions, each with a different distance to the closest choice color to the target, and trials were conducted in a random order. The trials were not speeded, and the colors remained visible until participants chose an option. To be conservative about the inclusion of participants, we excluded any participant who made an incorrect response in any of the 10 trials where the target color exactly matched one of the choice colors, leading to the exclusion of 7 of the 40 participants, and based on our a priori exclusion rule, excluded any participants whose overall accuracy was 2 standard deviations below the mean, leading to the exclusion of 0 additional participants. In addition, based on an a priori exclusion rule, we excluded trials with reaction times <200ms or >5000ms, which accounted for 1.75% (SEM:0.5%) of trials. The data from a subset of offset conditions is plotted in Figure 1C, and the full dataset is plot in Extended Data Fig. 1.

### 2. Psychophysical scaling triad experiment

N=100 participants on Mechanical Turk judged which of two colors presented was more similar to a target color, as in the fixed distance triad experiment. However, the pairs of colors now varied in offset from each other and from the target to allow us to accurately estimate the entire psychophysical distance function. In particular, the closest choice item to the target color could be one of 21 distances away from the target color: 0, 3, 5, 8, 10, 13, 15, 20, 25, 30, 35, 45, 55, 65, 75, 85, 100, 120, 140, 160, or 180 degrees. If we refer to these offsets as *o_i_*, such that o_1_ is 0 degrees offset and o_21_ is 180 degrees offset, then given a first choice item of o_i_, the second choice item was equally often o_i+1_, o_i+2_, o_i+3_, o_i+4_, or o_i+8_ degrees from the target color (excluding cases where such options were >21).

Participants were asked to make their judgments solely based on intuitive visual similarity and to repeat the word ‘the’ for the duration of the trial to prevent the usage of words or other verbal information. Each participant completed 261 trials, including 3 repeats of each of the possible pairs of offset conditions, and trials were conducted in a random order. The trials were not speeded, and the colors remained visible until participants chose an option. Following our a priori exclusion rule, we excluded any participant whose overall accuracy was 2 standard deviations below the mean (M=77.5%) leading to the exclusion of 8 of the 100 participants. In addition, based on an a priori exclusion rule, we excluded trials with reaction times <200ms or >5000ms, which accounted for 1.7% (SEM:0.26%) of trials.

To compute psychophysical similarity from these data, we used a modified version of the model proposed by Maloney and Yang^16^, the Maximum Likelihood Difference Scaling method. Rather than using this model to measure the distance between e.g., red and green, we adapted it to measure the appropriate psychophysical scaling of similarity between colors as a function of their distance between colors along the wheel rather than their absolute color. In particular, any given trial has a target color, S_i_, and two options for which is more similar, S_j_ and S_k_,. Let l_ij_ = S_j_ – S_i_, the distance between Si and Sj on the color wheel, which is always in the set [0, 3, 5,…180], and ψ_ij_, the psychophysical similarity to which this distance corresponds. If people made decisions without noise then they should pick item j if and only if ψ_ij_ > ψ_ik_. We add noise by assuming participants decisions are affected by Gaussian error, such that they pick item j if ψ_ij_ + ε > ψ_ik_. We set the standard deviation of the Gaussian ε noise to 1, consistent with a signal detection model. Thus, the model has 20 free parameters, corresponding to the similarity scaling values for each possible distance length (e.g., how similar a distance of 5 or 10 on the color wheel really is to participants), and then we fit the model using maximum likelihood search (*fmincon* in MATLAB). Thus, these scaled values for each interval length most accurately predict observers’ similarity judgments, in that equal intervals in the scaled space are discriminated with equal performance. Once the scaling is estimated, we normalize the psychophysical scaling parameters so that psychophysical similarity ranges from 0 to 1.

We did not test all possible pairings, but simply a subset (5 different offsets) because collecting more pairs does not improve the estimate of the psychophysical scaling function much, if at all, since the pairs we used ‘overlap’ enough without doing all of them. Each possible pairing provides an estimate of a ‘slope’ on the psychophysical similarity graph. For each pair, the relevant part of the x-axis is known, and people’s d’ at discriminating each pair (“which is closer? target+10 degrees or target+45 degrees”?) is an estimate of the y-axis difference / slope in that range (i.e. the difference in psychophysical similarity between those two points). Having 21 (distances) * 5 (offsets from those distances) = 105 such slope estimates, some covering wide ranges of the x-axis and some small ranges, and each well estimated, is sufficient to constrain the global shape of the function when using the MLDS method.

### 3. Likert color similarity experiment

N=50 participants on Mechanical Turk judged the similarity of two colors presented simultaneously on a Likert scale, ranging from 1 (least similar) to 7 (most similar). The colors were chosen from a wheel consisting of 360 color values that were evenly distributed along the response circle in the CIE *L*\**a*\**b*\* color space. The pairs of colors were chosen by first generating a random start color from the wheel and then choosing an offset (in degrees) to the second color, from the set 0, 5, 10, 20, 30, 40, 50, 60, 70, 80, 90, 120, 150, 180. Participants were given instructions by showing them two examples: (1) in example 1, the two colors were identical (0 deg apart on the color wheel) and participants were told they should give trials like this a 7; (2) in example 2, the two colors were maximally dissimilar (180 deg apart on the color wheel) and participants were told they should give this trial a 1. No information was given about how to treat intermediate trials. Participants were asked to make their judgments solely based on intuitive visual similarity and to repeat the word ‘the’ for the duration of the trial to prevent the usage of words or other verbal information. Each participant did 140 trials, including 10 repeats of each of the 14 offset conditions, each with a different starting color, and trials were conducted in a random order. The trials were not speeded, and the colors remained visible until participants chose an option. 2 participants were excluded for failing a manipulation check (requiring similarity >6 for trials where the colors were identical). Based on an a priori exclusion rule, we excluded trials with reaction times <200ms or >5000ms, which accounted for 3.0% (SEM:0.4%) of trials. Similarity between two colors separated by *x*° was measured using a 7-point Likert scale, where *S_min_* = 1 and *S_max_* = 7. To generate the psychophysical similarity function, we simply normalize this data to range from 0 to 1, giving a psychophysical similarity metric, such that *f*(*x*) = ((*S_x_* - *S_min_*) / (*S_max_* - *S_min_*)).

### 4. Perceptual matching experiment

N=40 participants on Mechanical Turk were shown a color and had to match this color, either using a continuous report color wheel (100 trials) or choosing among 60 options (100 trials; spaced 6 degrees apart on the color wheel, always including the target color). The 60-AFC version was designed to limit the contribution of motor noise, since the colors in this condition were spaced apart and presented as discrete boxes that could not easily be ‘misclicked’. Colors were generated using the same color wheel as other experiments, and participants had unlimited time had to choose the matching color. The color and color wheel/response options remained continuously visible until participants clicked to lock in their answer. The color was presented at one of 4 locations centered around fixation (randomly), approximately matching the distance to the color wheel and variation in position used in the continuous report memory experiments. 1 participant’s data was lost due to experimenter error and 2 participants were excluded for an average error rate greater than 2 standard deviations away from the mean.

To convert this data into a perceptual correlation matrix —asking how likely participants are to confuse a color *x* degrees away in a perception experiment — we relied upon the 60-AFC data alone, since this data has no contribution from motor noise and so is solely a measure of perceptual noise. However, using the continuous report data instead result in no difference in any subsequent conclusions, as the contribution of motor noise in that task appeared to be minimal. To create the perceptual correlation matrix, we created a normalized histogram across all participants of how often they made errors of each size up to 60 degree errors (−60, −54… 0,… 54, 60), and then linearly interpolated between these to get a value of the confusability for each degree of distance. We then normalized this to range from 0 to 1.

### 5. Modeling Data Using the Target Confusability Competition (TCC) Model

The model is explained interactively here: https://bradylab.ucsd.edu/tcc/ In general, the model is typical of a signal detection model of long-term memory, but adapted to the case of continuous report, which we treat as a 360 alternative forced-choice for the purposes of the model. The analysis of such data focuses on the distribution of errors people make measured in degrees along the response wheel, *x*, where correct responses have *x*=0° error, and errors range up to *x*=±180°, reflecting the incorrect choice of the most distant item from the target on the response wheel (Fig. 1B). In the TCC model, when probed on a single item and asked to report its color, (1) each of the colors on the color wheel generates a memory-match signal *m_x_*, with the strength of this signal drawn from a Gaussian distribution, *m_x_* ~ *N*(*d_x_*, 1), (2) participants report whichever color x has the maximum *m_x_*, (3) the mean of the memory-match signal for each color, *d_x_*, is determined by its psychophysical similarity to the target according to the measured function (*f*(*x*)), such that *d_x_* = *d*′ *f*(*x*) (Figure 2) and (4) the noise is correlated across nearby colors according to confusability in a perceptual matching task. As *f*(*x*), the psychophysical similarity function, we use the smooth function estimated from the Likert similarity experiment although the triad task modeled similarity function predicts fundamentally the same results (Extended Data Fig. 4).

According to the model, the mean memory-match signal for a given color *x* on the working memory task is given by *d_x_* = *d*′ *f*(*x*), where *d*′ is the model’s only free parameter. When *x* = 0, *f*(*x*) = 1, so *d*_0_ = *d*′. By contrast, when *x* = 180, *f*(*x*) = 0, so *d_180_* = 0. Then, as noted above, at test each color on the wheel generates a memory-match signal, *m_x_*, conceptualized as a random draw from that color’s distribution centered on *d_x_*. That is, if the noise was uncorrelated between nearby colors, *m_x_* ~ *N*(*d_x_*, 1). The response (*r*) on a given trial is made to the color on the wheel that generates the maximum memory-match signal, *r* = *argmax*(*m*).

Thus, the full code for sampling an absolute value of error from such a TCC-like (uncorrelated noise) model is only two lines of MATLAB:

~~~
memMatchStrengths = randn(1,180)+
similarityFunction * dprime;
[~,memoryError] = max(memMatchStrengths);
~~~

This model fits the data well as-is (see Extended Data Fig. 2), but as specified so far, this model assumes that 360 independent color probes elicit independent noisy memory-match signals. The shape of the distributions the model predicts are effectively independent of how many color channels we assume, so this number is not important to TCC’s ability to fit working memory data, but the *d*′ value in the model does change depending on the number of color channels used. Thus, to make the *d*′ value in TCC comparable to real signal detection *d*′ values, it is important to consider “how many” color channels people are accessing.

Rather than make this a discrete decision (e.g., ‘there are 30 independent colors on the color wheel, so people consider 30 channels’), we instead estimated the covariance between nearby channels in a continuous manner. The familiarity value of color 181 and 182 on the wheel cannot possibly be fully independent, since these two colors are perceptually indistinguishable. Following this intuition, we make a simple assumption: the amount of shared variance in the noise between any two color channels is simply how often colors at that distance are confused in a perceptual matching task. Thus, *p*(*x*), the correlation in the noise between any two colors *x* apart on the color wheel, is given by C_x_/ C_0_, where C_x_ is how often colors *x* degrees away from the target are chosen in the perceptual matching task (with these values interpolated from the histogram of errors; see Methods section 4). On average, colors 1 degree away are chosen about 96% as often as the correct color in the matching task, so the noise between any two channels 1 degree apart is assumed to share 96% of its variance; 82% at 5 degrees; etc. Thus, having measured both the similarity function and the perceptual matching matrix, to sample from the full (correlated-noise) TCC model, we can use MATLAB code that is nearly as straightforward as the uncorrelated model:

~~~
memMatchStrengths = mvnrnd(similarityFunction *
dprime, percepCorrMatrix);
[~,memoryError] = max(memMatchStrengths);
~~~

Thus, in the full version of TCC, the mean of the memory-match signal for each color, *d_x_*, is determined by its psychophysical similarity to the target according to the measured function *f*(*x*), which is taken to be symmetrical for the fitting based on the averaged similarity data, such that *d_x_* = *d*′ *f*(|*x*|), for *x*’s [−179,180]. The covariance between colors (*R*) is given by the perceptual confusability of colors at that distance, *p*(*x*), which is also taken to be symmetric:

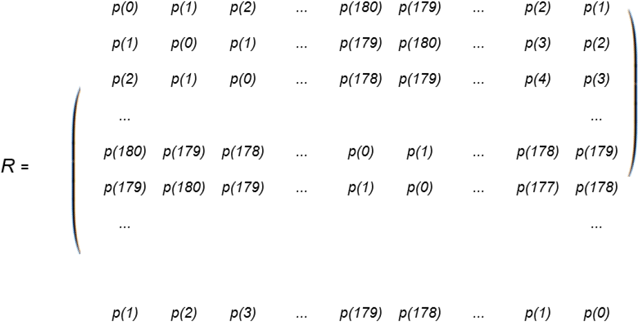

To use the perceptual correlation data as the covariance in the correlated model, because it might not always be a perfect correlation matrix (e.g., not perfectly symmetric, as it was based on real data), we first computed *R* and then iteratively removed negative eigenvalues from this matrix and forced it to be symmetric until it was a valid correlation matrix. This resulted in only minimal changes compared to the raw perceptual correlations inferred from the perceptual confusability data.

Then let (X_-179_,…, X_180_) be a multivariate normal random vector with mean *d*, unit variance, and correlation matrix *R*. The winning memory strength (*m*; i.e., subjective confidence) and reported color value, *r*, are then the max and argmax, respectively, of this vector:

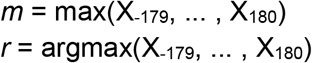

And the error, *e*, is the circular distance from *r* to *0*. The distribution of *m* is in theory directly computable^56^, but we rely on sampling from this distribution for the fits in the current paper (see below).

Although also not important to the fit of the current data, the model can also be adapted to include a motor error component. Whereas existing mixture models predict the shape of the response distribution directly and thus confound motor error with the standard deviation of memory (see Fougnie et al.^57^ for an attempt to de-confound these), our model makes predictions about the actual item that participants wish to report. Thus, if participants do not perfectly pick the exact location of their intended response on a continuous wheel during every trial, a small degree of Gaussian motor error can be assumed to be included in responses, e.g., the response on a given trial, rather than being argmax(X_-179_,…, X_180_), likely includes motor noise of some small amount, for example, 2°:

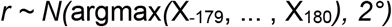

Thus, for accuracy to the real generative model of responses, in the model fitting reported in the present paper, we include a fixed normally distributed motor error with SD=2°, although we found the results are not importantly different if we do not include this in the model.

For fits using the uncorrelated noise model, fits of the *d*’ parameter of the model to datasets were performed using the MemToolbox^58^ making use of maximum likelihood (see code on OSF). For fits of the correlated model —which is difficult to compute a likelihood function for but straightforward to sample from —we relied on sampling 500,000 samples from the model’s error at each of a range of d’ values (0 to 4.5 in steps of 0.02) and slightly smoothing the result to get a pdf for the model at each *d*′ value. The uncorrelated noise version of TCC —which can be directly maximized —results in the same fits as the correlated version, with *d*′ linearly scaled by ~0.65. (See Extended Data Figure 2). Thus, it is also possible to fit the correlated noise version by fitting the uncorrelated version through maximum likelihood with the appropriate adjustment to *d*′, and doing so results in the same fits.

### 6. Continuous color report data (set size 1, 3 and 6, 8)

The continuous color report data used for fitting the model was collected in the lab to allow a larger number of trials per participant. N = 20 participants performed 100 trials of a memory experiment at each of set size 1, 3, 6 and 8, for a total of 400 trials (plus 4 practice trials). The display consisted of 8 placeholder circles. Colors were then presented for 1000ms, followed by an 800ms ISI. For set sizes below 8, the colors appeared at random locations with placeholders in place for any remaining locations (e.g. at set size 3, the colors appeared at 3 random locations with placeholders remaining in the other 5 locations). Colors were constrained to be at least 15° apart in color space along the response wheel. After the ISI, a target item was probed by marking a placeholder circle was marked with a thicker outline, and participants were asked to respond on a continuous color wheel to indicate what color had been presented at that location. The response wheel was held constant from trial-to-trial. Error was calculated as the number of degrees on the color wheel between the probed item and the response. No subjects were excluded.

### 7. Continuous report memory as a function of delay (set size 1, 3, 6)

N = 20 participants in-lab completed a color working memory task similar to the previous high set size experiment, but with the following exceptions. Participants performed 12 blocks of 75 trials (900 trials total). Each block contained an equal number of trials at set size 1, 3, and 6. The display consisted of 6 placeholder circles. Colors were presented for 500ms, and followed by a delay of either 1000ms, 3000ms, or 5000ms. Delay time was blocked, and participants were informed at the beginning of each block the delay time for that block. Each combination of the 3 set sizes and the 3 delays was used in 100 trials. One participant was excluded for having performance greater than 2 standard deviations worse than average (across all conditions), leaving a final sample of 19.

### 8. Continuous report memory as a function of encoding time (set size 1, 3, 6)

N = 20 participants in-lab completed a color working memory task identical to the delay experiment, but with the following exceptions. Participants performed 12 blocks of 75 trials. Each block contained an equal number of trials at set size 1, 3, and 6. Colors were presented for either 100ms, 500ms, or 1500ms. Encoding time was blocked, and participants were informed at the beginning of each block the encoding time for that block. Following encoding, there was a 1000ms delay before a target item was probed. Each combination of the 3 set sizes and the 3 encoding times was used in 100 trials. No subjects were excluded.

### 9. Model comparisons to mixture models

For all model comparisons in the paper, we created new versions of mixture models designed to be directly comparable to TCC. In particular, to make predictions derived from mixture models comparable to those derived from TCC (which specifies a probability of response discretely for each 1 degree of the wheel, not over a continuous distribution), we use discrete versions of the 2-parameter and 3-parameter mixture models in which the probabilities of the data are normalized over each of 360 possible integer error values (not over the continuous space of errors).

We performed two types of model comparisons: one to simply assess the fit of the model to the data, and one designed to penalize more complex models. In particular, we first performed a cross-validation procedure to assess the fit of each model^59^. Specifically, we fit the TCC and the 2-parameter and variable precision mixture models to data from each set of N-1 trials separately for each subject and set size and then evaluated the log-likelihood of this model using data from the single held out trial. We then assessed the reliability of this likelihood difference across subjects separately for each set size. TCC and mixture models provided relatively comparable fits (see Supplementary Table 2), which is to be expected given the mixture model can almost perfectly accurately mimic TCC (see Supplementary Fig. 3) and given that the amount of data used to fit the models is much greater than the number of parameters in either model (which ranges from 1-3), so cross-validation provides effectively no penalty for complexity.

We then compared how well the competing models (TCC; 2-parameter mixture model; 3-parameter variable precision mixture model) fit data from individual participants for the color report data when using an explicit penalty for the greater complexity of the mixture models. In particular, we assessed BIC separately for each set size and each participant. We found a strong preference for TCC over both mixture models when penalizing complexity (see Supplementary Table 2). Note that this was true even though TCC fits are based on aggregated similarity functions from a different group of participants, collected in a different way (online vs. in lab), suggesting the global structure of the psychophysical similarity function is largely a fixed aspect of a given stimulus space. Ideally, TCC would be fit with a similarity function specific to each individual target color (which can be done and predicts the appropriate deviations; see Extended Data Fig. 5), which would almost certainly improve the fit of TCC even further with no added parameters (because the added complexity would simply be more measured perceptual data). However, in the current fits we rely solely on averaged similarity to demonstrate how it is the global, not local, structure of the similarity space that is critical to the fit of TCC.

### 10. 2-AFC at different foil similarities

N=60 participants on Mechanical Turk completed 200 trials of a 4-item working memory task. On each trial, they saw 4 colors randomly chosen from the color wheel (subject to the constraint that no two colors were within 15 deg. of each other). The colors were presented for 1000ms and then after an 800ms delay, had to answer a 2-AFC memory probe about one of the colors. The foil color in the 2-AFC could be offset from the target 180, 72, 24, or 12 degrees (50 trials/condition). These conditions were interleaved so that participants needed to maintain detailed memories of the color on every trial, since conceivably if only 180 degree foils were present for a block or in an entire experiment, participants would be likely to encode only categorical, not perceptual information. The response options were presented at appropriate locations along a full color wheel -- e.g., the 180 degree foils were presented 180 deg. apart on the screen, and the 12 deg. foils were presented 12 deg. apart on the screen, to visually indicate the distance between the target and foil in color space. The response wheel was rotated from trial-to-trial.

Performance was scored as the number correct out of 50 at each offset of the memory foil. 5 participants were excluded for below chance performance in the maximally easy 180 deg. offset condition, leaving N=55 participants.

In order to assess the predictions of TCC for this data in a way amenable to the use of Bayes factors, we took the number correct out of 50 in the 180 deg. foil condition and used this to calculate a probability distribution over d’ values (e.g., any given *d*′ predicts, according to the binomial function, a likelihood over all numbers of correct responses). In TCC, a given *d*′ value for 180 deg. foils predicts *d*′ for all other offsets straightforwardly, although for the correlated-noise TCC, performance is not simply *d*′ modulated by similarity (for similar foils, the correlated noise plays a role). Thus, to predict performance we sampled from the model repeatedly, e.g., for 24 deg. foils, in MATLAB notation:

~~~
memoryMatchStrengths =
mvnrnd(similarityFunction * dprime_180_,
percepCorrMatrix, 50);
isCorrect=memoryMatchStrengths_0deg_>memoryMatchStr
engths_24deg_
~~~

In other words, to assess performance in the 24 deg. offset condition, we assumed responses were generated according to the argmax of only these values:

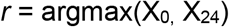

To preserve all uncertainty, we marginalized over the distribution of *d*′ values implied by the number of correct trials in the 180 deg. foil case and used this to make a prediction about the distributions of correct answers expected for each of the other offset conditions. This allows us to understand the likelihood of each subjects’ performance in the other conditions given their 180 deg. foil performance in TCC.

To assess the likelihood of performance at different offsets in the mixture model framework of Zhang and Luck^7^, we use performance at the 180 deg. foil conditions to assess the “guess rate” of participants (guess rate = 1 - (2*percentCorrect_180_-1)) in the standard way (e.g., Brady et al.^60^). However, in this framework, 180 deg. foils leave an unknown free parameter: memory precision cannot be assessed using such foils, and thus is free to vary. Thus, to predict a likelihood of each performance level at each other foil offset, we needed to marginalize over the unknown precision parameter. To minimize assumptions about this, we used the same prior on precisions that van den Berg et al.^8^ used when fitting both the standard mixture model and their own variable precision model, a uniform prior over the concentration parameter of the von Mises from 0-200. For any given ‘guess rate’ and ‘precision’, we then calculated the percentage of the PDF that was closest to each 2-AFC response option at each offset to generate a likelihood for the data (as in MemToolbox^58^). To calculate Bayes factors, we used a grid of values for both the *d*′ in TCC and for the precision in the mixture model, using steps of 1 in the precision and steps of 0.01 in *d*′, and we assessed the summed log likelihood of each of the 3 other offsets (e.g., not including the 180 deg. condition) as our final data likelihood.

### 11. 2-AFC generalization to n-AFC and continuous report

N=60 participants on Mechanical Turk completed 200 trials of a 4-item working memory task. On each trial, they saw 4 colors randomly chosen from the color wheel (subject to the constraint that no two colors were within 15 deg. of each other). The colors were presented for 1000ms and then after an 800ms delay, had to answer a probe about one of the colors. This probe could be a 2-AFC (with 180 deg. different foil), an 8-AFC (with the choices equally spaced around the color wheel, and always including the target), a 60-AFC (similarly equally spaced), or continuous report (360-AFC). These conditions were interleaved so that participants needed to maintain detailed memories of the color on every trial, since conceivably if only 180 degree foils were present for a block or in an entire experiment, participants would be likely to encode only categorical, not perceptual information. The response options were presented at appropriate locations along a full color wheel -- e.g., the 2-AFC 180 degree foils were presented 180 deg. apart on the screen, and the 60-AFC foils deg. foils were presented 6 deg. apart on the screen, to visually indicate the distance between the target and foils in color space. The response wheel was rotated from trial-to-trial.

Performance was scored as the number correct out of 50 at each offset of the memory foil. One participant’s data was lost, and 7 participants were excluded for below chance performance in the maximally easy 2-AFC, 180 deg. offset condition, leaving N=52 participants.

The simplest metric is simply to compare the d’ computed from 2-AFC performance (e.g., (norminv(hit)-norminv(fa))/sqrt(2)) to the d’ from fitting TCC to the continuous report data. These are extremely strongly related (Fig. 5B).

In order to assess the predictions of TCC for this data in a way amenable to the use of Bayes factors, we again took the number correct out of 50 in the 2-AFC 180 deg. foil condition and used this to calculate a distribution over d’ values (e.g., any given *d*′ predicts, according to the binomial function, a likelihood over all numbers of correct responses). In TCC, a given *d*′ value for 180-foils predicts *d*′ for all other n-AFCs (including 360-AFC) straightforwardly, by simply first choosing the maximum out of the relevant foil options that are present, e.g., at 8-AFC:

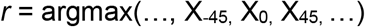

To preserve all uncertainty, we marginalized over the distribution of *d*′ values implied by the number correct in the 180 deg. foil case and used this to make a prediction about the distributions of responses to each foil expected for each of the other n-AFC conditions. This allows us to understand the likelihood of each subjects’ performance in the other conditions given their 180 deg. foil performance in TCC.

To assess the likelihood of performance in continuous report given performance in the 2-AFC task, in the mixture model framework of Zhang and Luck^7^, we use performance at the 180 deg. foil conditions to assess the “guess rate” of participants (guess rate = 1 - (2*percentCorrect_180_-1)) in the standard way (e.g., Brady et al.^60^). However, in this framework, 180 deg. foils again leave an unknown free parameter: memory precision cannot be assessed using such foils, and thus is free to vary. Thus, to predict a likelihood of each performance level at each other foil offset, we needed to marginalize over the unknown precision parameter. To minimize assumptions about this, we used the same prior on precisions that van den Berg et al.^8^ used when fitting both the standard mixture model and their own variable precision model, a uniform prior over the concentration parameter of the von Mises from 0-200. For any given ‘guess rate’ and ‘precision’, we then calculated the likelihood of subject’s continuous report performance under these parameters. To calculate Bayes factors, we used a grid of values for both the *d*′ in TCC and for the precision in the mixture model, using steps of 1 in the precision and steps of 0.01 in *d*′. We assessed the log likelihood of TCC and the mixture model only in the continuous report case, having fit the parameter(s) using only the data from the 2-AFC 180 deg. condition.

### 12. Face identity continuous report data (set size 1 and 3)

We utilized the same continuous report task, but adapted the stimulus space to face identity using the continuous face identity space and continuous response wheel created in Haberman, Brady and Alvarez^29^. In particular, as described in that work, the faces were 360 linearly interpolated identity morphs, taken from the Harvard Face Database, of three distinct male faces (A-B-C-A; see Figure 6), generated using MorphAge software(version 4.1.3, Creaceed). Face morphs were nominally separated from one another in identity units, which corresponded to steps in the morph space. Prior to morphing, face images were luminance normalized. In our memory task, we used set sizes 1 and 3, showing either 1 or 3 faces at once, and the encoding display was shown for 1.5 seconds due to the increased complexity of the face stimuli and task difficulty. Participants on Mechanical Turk (N=50) completed 180 trials. The first 20 trials were practice and not included in the analysis. 14 participants were excluded for having near-chance performance levels (*d*′<0.50) at set size 3, although including all subjects with *d*′>=0 does not affect our conclusions or the fit of TCC.

### 13. Face identity similarity ‘quad’ task

N=102 participants on Mechanical Turk judged which of two pairs of faces presented were more distinct (e.g., which pair had constituent items that were more different from each other). On each trial, we chose 2 pairs of faces, with the first item in each pair being randomly chosen and the second item in each pair always having an offset of 0, 5, 10, 20, 40, 60, 80, 100, 140, or 180 away. Altogether, they completed 18 trials of each kind, giving a total of 180 trials each.

Participants were asked to make their judgments solely based on intuitive visual similarity, rather than the use of knowledge of faces or using verbal labels. We excluded participants whose overall performance level was more than 2 standard deviations below the mean, resulting in a final sample of N=85.

To compute psychophysical distance from these data, we used a similar model as for colors, based on the model proposed by Maloney and Yang^16^, the Maximum Likelihood Difference Scaling method. In particular, any given trial has two pairs of faces, where their face-wheel values are, S_i_, S_j_ and S_k_, S_l_. Let l_ij_ = S_j_ – S_i_, the length of the physical interval between S_i_ and S_j_, which is always in the set [0, 5, 10…180], and ψ_ij_, the psychophysical similarity to which this distance corresponds. If people made decisions without noise then they should pick pair i,j if and only if ψ_ij_ > ψ_kl_. We add noise by assuming participants decisions are affected by Gaussian error, such that they pick pair i,j if ψ_ij_ + ε > ψ_kl_. We set the standard deviation of the Gaussian ε noise to 1, so that the model has 9 free parameters, corresponding to the psychophysical scaling values for each possible interval length (e.g., how similar a distance of 5 or 10 ‘really’ is to participants), and then we fit the model using maximum likelihood search (*fmincon* in MATLAB). Thus, these scaled values for each interval length most accurately predict observers’ judgments, in that equal intervals in the scaled space are discriminated with equal performance. Once the scaling is estimated, we normalize the psychophysical scaling parameters so that psychophysical similarity ranges from 0 to 1.

### 14. Face identity perceptual matching

N=40 participants on Mechanical Turk were shown a face and had to match this face using a continuous report wheel (100 trials). Because the contribution of motor noise appeared to be minimal in the color matching task (relative to perceptual error), we used only a continuous report wheel (no 60-AFC). Faces were generated from the same continuous face space used in other experiments, and participants had unlimited time had to choose the matching face. The face and face wheel/response options remained continuously visible until participants clicked to lock in their answer. The face was presented at one of 4 locations centered around fixation (randomly), approximately matching the distance to the face wheel and variation in position used in the continuous report memory experiments. 7 participants were excluded for below chance error rates.

To convert this data into a perceptual correlation matrix -- asking how likely participants are to confuse a face *x* degrees away in a perception experiment -- we created a normalized histogram across all participants of how often they made errors of each size (in bins of 5 deg.: −180, −175,… 180) and then linearly interpolated between these to get a value of the confusability for each degree of distance. We then normalized this to range from 0 to 1.

### 15. Visual long-term memory color report task

Long-term memory data from Fig. 6 was taken from Miner, Schurgin, Brady^30^, Experiment 2A. N=30 participants in the lab at UC San Diego performed 5 blocks of a long-term memory experiment. In each block they memorized real-world objects’ colors, and then after a brief delay, were shown a sequence of memory tests. Each block’s study session consisted of 20 items of distinct categories seen only once and 10 items also of distinct categories seen twice, for a total of 40 presentations of colored objects. Each presentation lasted 3 seconds followed by a 1 second inter-stimulus interval. At test, 20 old objects were presented (10 seen once, 10 seen twice) and 20 new objects of distinct categories were presented. Participants saw each object in grayscale and made an old/new judgment, and then, if they reported the item was old, they reported its color using a continuous color wheel. As described in Miner et al.^30^, 6 participants were excluded per the criterion used in that paper.

Note that long-term memory performance in this task likely depends on a two part decision -- item memory and source memory (e.g., the object itself, and then its color). This two-part decision is related to the processes of recollection and familiarity that can be modeled in various ways^61^, and likely introduces significant heterogeneity into the color memory strength, since some items will have weak item memories, preventing the retrieval of color information. TCC provides a strong fit here, and to the other long-term memory data plotted in Fig. 8, without addressing this, likely due to the fact that item memory in all of these studies was very strong (only a small number of categorically distinct items needed to be remembered). Future research should clarify how TCC connects to distinctions between recollection and familiarity and the extent to which heterogeneity in d’ between items in long-term memory must be assumed for fitting a wider variety of tasks.

### 16. Literature Analysis

To assess our model’s prediction that previously observed trade-offs between different psychological states are measuring the same underlying parameter (*d*′), we conducted a literature analysis of data from color working memory research. In particular, we examined the two parameters most commonly reported by those fitting mixture models to their data, precision (in terms of SD) and guessing.

We searched for papers in mid-2018 that used these mixture model techniques by finding papers that cited the most prominent mixture modeling toolboxes, Suchow, Brady, Fougnie & Alvarez^58^ and Bays et al.^62^. We used a liberal inclusion criteria in order to obtain as many data points as possible. Our inclusion criteria were papers that cite either of these toolboxes and report data where: 1) There was some delay between the working memory study array and test; 2) Instructions were to remember all the items; 3) SD/guess values were reported or graph axes were clearly labeled; 4) Participants were normal, healthy, and between ages 18-35. 5). Colors used were widely spaced, discriminable colors from the CIE *L*\**a*\**b*\* color space. Note that even slight changes in the color wheel used between papers (or the addition of noise to stimuli^7^) changes the perceptual confusability of the stimuli and therefore ideally calls for a different similarity function to be measured and therefore a different prediction from TCC about the relationship between ‘guess rate’ and ‘SD’. However, in the current literature analysis we simply assumed these were the same for all papers. For papers that did not report SD/guess values in the text or tables, these values were obtained by digitizing figures with clear axis labels^63^.

These inclusion criteria resulted in a diverse set of data points, including studies using sequential or simultaneous presentation, feedback vs no feedback, cues vs no-cues, varying encoding time (100-2000 ms), and variable delay (1-10 sec). A total of 14 papers and 56 data points were included (Supplementary Table 4). In general, TCC provides a strong fit to this existing data given the heterogeneity in methods (Supplementary Fig. 4) and this data is also consistent with the idea that there is no added ‘guessing’ at high set sizes (Supplementary Fig. 5).

## Data Availability Statement

All relevant data for this manuscript are available at: https://osf.io/j2h65/?view_only=fdd51dd775a945508c7cbbf25b662692

## Code Availability Statement

All relevant analysis code for this manuscript is available at: https://osf.io/j2h65/?view_only=fdd51dd775a945508c7cbbf25b662692

## Acknowledgements

We thank Viola Störmer, Rosanne Rademaker, Jeremy Wolfe, Maria Robinson, Daryl Fougnie and Talia Konkle for comments on these ideas and on the manuscript, and Yong Hoon Chung and Brittany Hawkins for help with data collection. For funding, we would like to acknowledge NSF CAREER (BCS-1653457) to TFB. The funders had no role in study design, data collection and analysis, decision to publish or preparation of the manuscript.

## Author Contributions

M.W.S. jointly conceived of the model with J.T.W. and T.F.B; M.W.S. and T.F.B. designed the experiments; T.F.B. wrote code, ran the model, and analyzed output data; M.W.S., J.T.W., and T.F.B. wrote the manuscript.

## Extended Data Figures

**Extended Data Figure 1.**
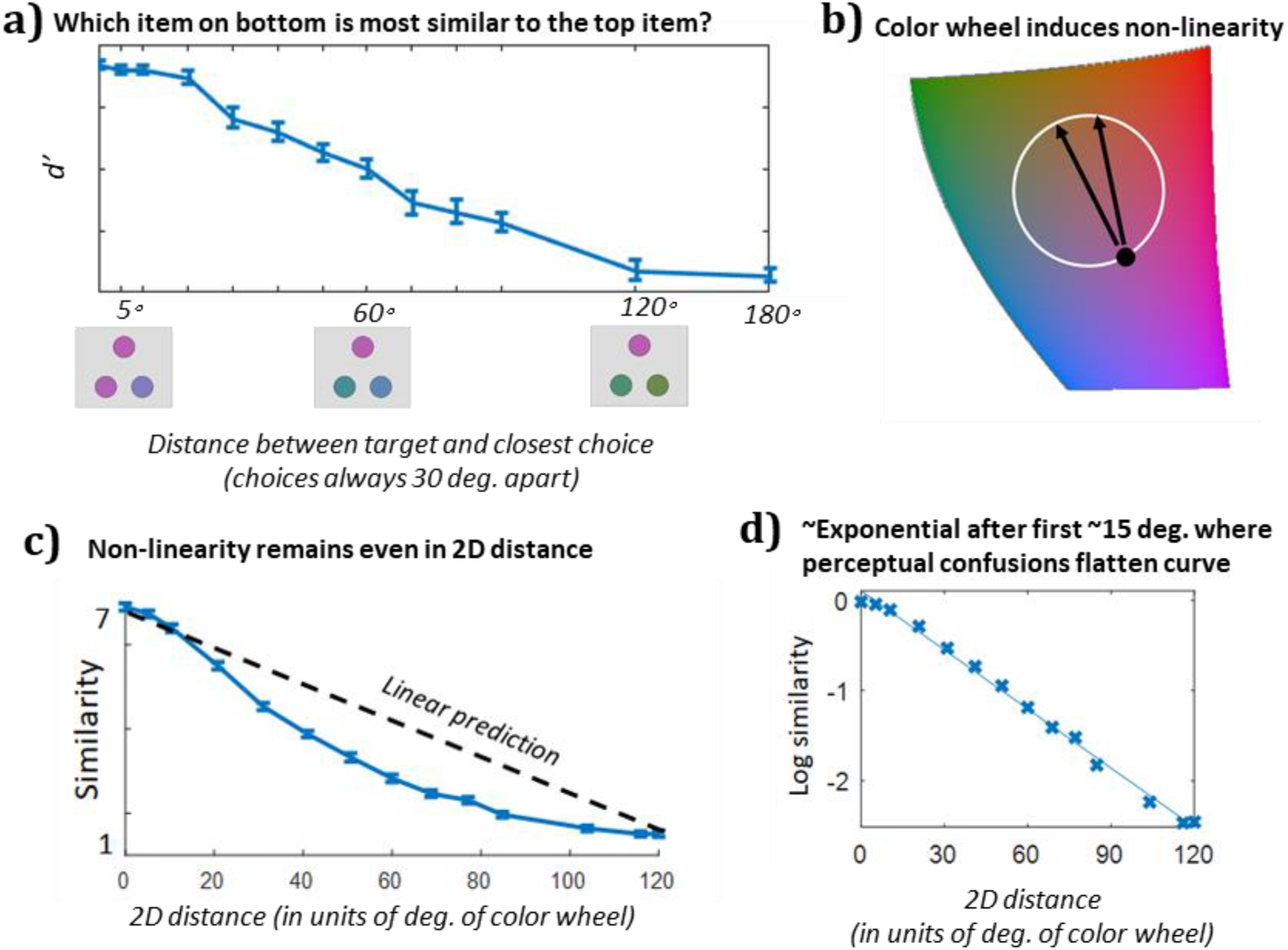
**(A)** Data from all distances in the fixed distance triad task (Figure 1C). On each trial, there was a target color, here always at 0°, and participants’ task was to choose which of two other colors was closer to the target color in color space. The two choice colors always differed by 30°. The x-axis shows the closer color of the two choice colors. That is, the 150° label on the x-axis reflects performance on a condition where the two choices were 150° and 180° away from the target color. As shown with a subset of this data in Figure 1C, increasing distance from the target results in a decreased ability to tell which of two colors is closer to the target in color space. This shows the non-linearity of color space with respect to judgments of color similarity. Note that this function does not depict the actual psychophysical similarity function: Roughly speaking, the *d*′ estimate in this graph is the estimate of instantaneous slope (over a 30 deg. range) in the similarity function in Figure 1F. (**B**) Despite being conceived of as a color wheel in many memory experiments, in reality, participants internal representation of color -- and thus the confusability between colors -- ought to be a function of their linear distance in an approximately 3D color space, not their angular distance along the circumference of an artificially imposed wheel. Since the colors are equal luminance, we can conceive of this on a 2D plane. Thus, on this plane the confusability of a color “180 degrees away” on the wheel is only slightly lower than one “150 degrees away” on the wheel, since in 2D color space it is only slightly further away. This simple non-linearity from ignoring the global structure of the color ‘wheel’ partially explains the long tails observed in typical color report experiments, although it does not explain the full degree of this non-linearity, which is additionally attributable to psychophysical similarity being a non-linear function even of distance across 2D color space. (**C**) The similarity function remains non-linear even in 2D color space. Distances here are scaled relative to the color wheel rather than in absolute CIELa*b* values., e.g., an item 180 degrees opposite on the color wheel is “120” in real distance since if the distance along the circumference is 180, 120 is the distance across the color wheel. (**D**) Plotted on a log axis, the similarity falls off approximately linearly, indicating that similarity falls of roughly exponentially with the exception of colors nearby the target. The non-exponential fall-off near the 0 point reflects perceptual noise/lack of perceptual discriminability between nearby colors. As shown in Figure 1, when you convolve measured perceptual noise with an exponential function, this provides a very good fit to the similarity function, consistent with a wide-variety of evidence about the structure of similarity and generalization^19^.

**Extended Data Figure 2.**
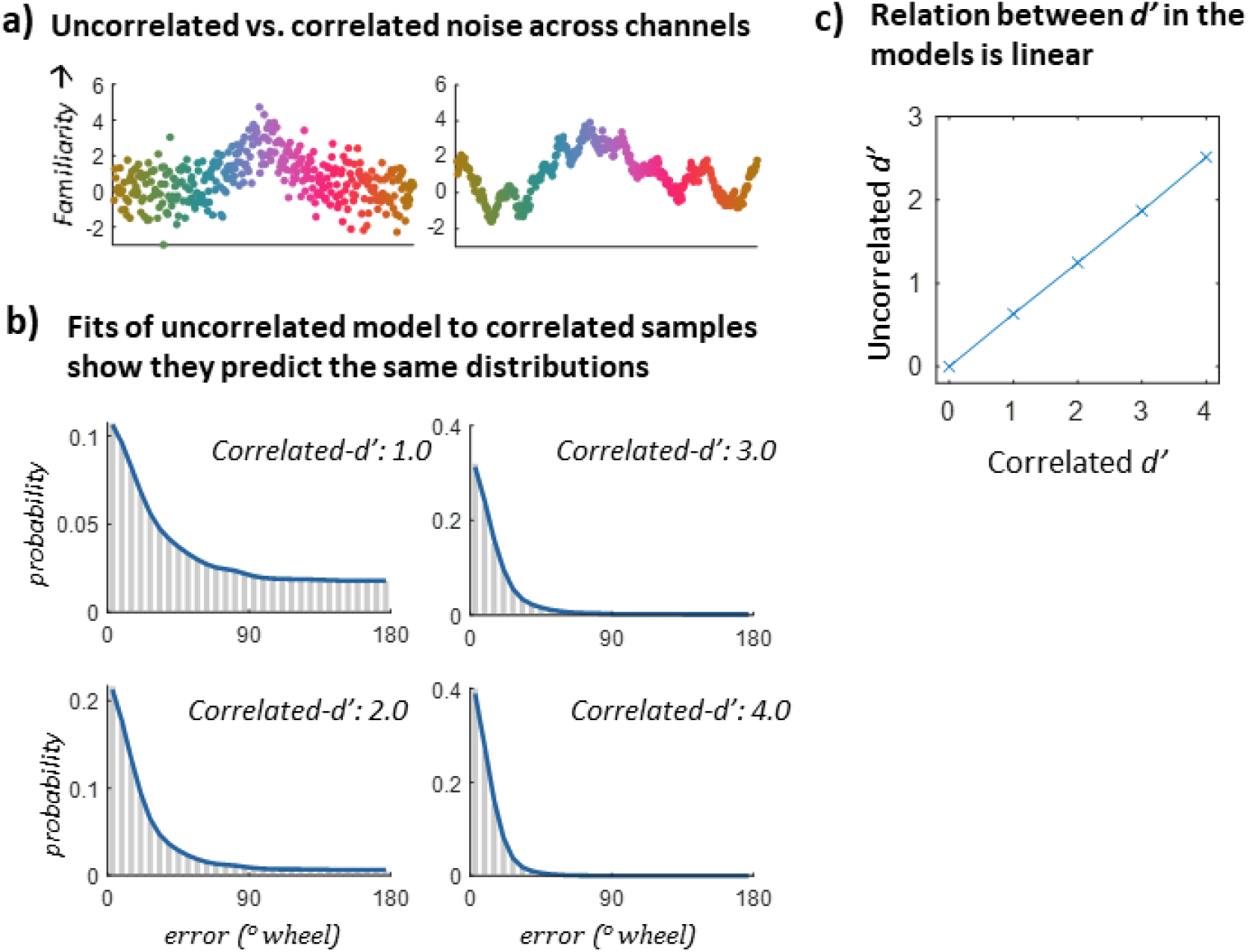
Simulations of uncorrelated vs. correlated noise versions of TCC. Only the correlated-noise TCC produces true *d*′ values -- those that are interchangeable with *d*′ you’d estimate from a same/diff task with the same stimuli. However, the simpler uncorrelated noise TCC predicts the exact same distributions of errors in continuous report, and the d’ values between the correlated and uncorrelated noise models are linearly related by a factor of ~0.65. Thus, in many cases it may be useful to fit the uncorrelated TCC to data and then adjust the *d*′ rather than fitting correlated noise TCC. This also means that for color, similarity alone without perceptual confusion data can be used to make linear (but not exact) predictions about confusability in n-AFC tasks outside the range of perceptual confusion (approx. 15 deg).

**Extended Data Figure 3.**
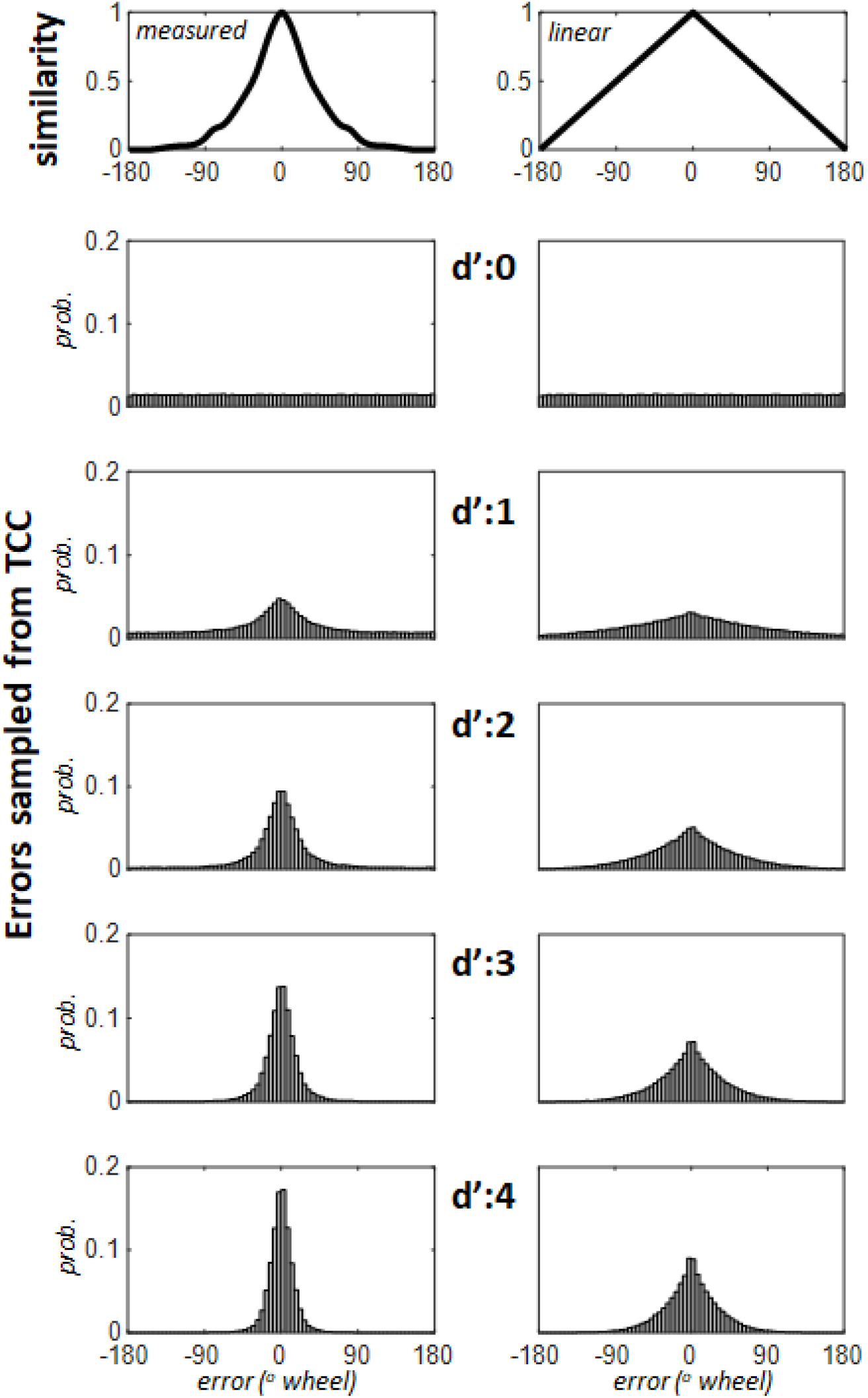
Simulations show data sampled from TCC, using either the measured psychological similarity function or a linear similarity function. Given a linear similarity function, it is clear TCC does not predict response distributions similar to human performance -- such fits are critically dependent on the well-known exponential-like shape of similarity functions. Notice also how the max rule from the signal detection decision process plays a major role in the shape of the distributions. Since people pick the strongest signal, the distribution of max signals is peakier than the underlying signals themselves (which always follows the similarity function).

**Extended Data Figure 4.**
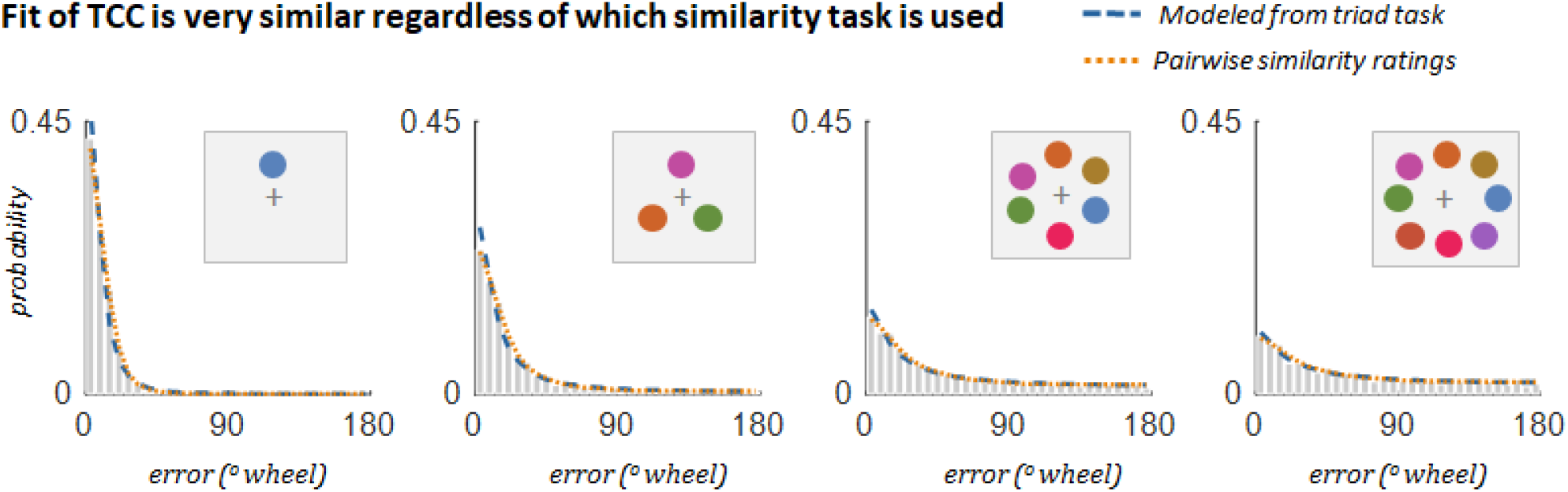
Comparison of fit to memory data for similarity functions reported in main text. In the current data for color, both the model-based triad psychophysical scaling data and the Likert similarity rating produce extremely similar data (see Figure 1). Thus, they all produce similar fits to the memory data (shown here are the set size data). It is important to note that depending on the number of trials, a large number of data points (i.e. subjects) may be necessary in order to obtain reliable estimates of a given stimulus space in the triad and quad scaling tasks (we use the quad task for face similarity). The Likert task requires considerably less data to estimate, and it was in agreement with the results of the triad task for colors, so we rely on it as our primary measure of similarity in the current fits. However, depending on the stimulus space, observers may utilize different strategies in such subjective similarity tasks (particularly for spaces, like orientation, where it is obviously a linear physical manipulation), and ultimately an objective task like the quad task may be best to understand the psychophysical similarity function. This is why for the face space task we used the quad similarity task. The task used to estimate similarity is important in that it is important that participants provide judgments of the absolute interval between stimuli and not rely on categories or verbal labels, or, in the triad task, that participants not rely on a relational or relative encoding of the two choice items rather than their absolute distance to the target item. How best to ensure that participants rely on absolute intervals is represented in a large literature dating to Thurstone^65^ and Torgerson^15^.

**Extended Data Figure 5.**
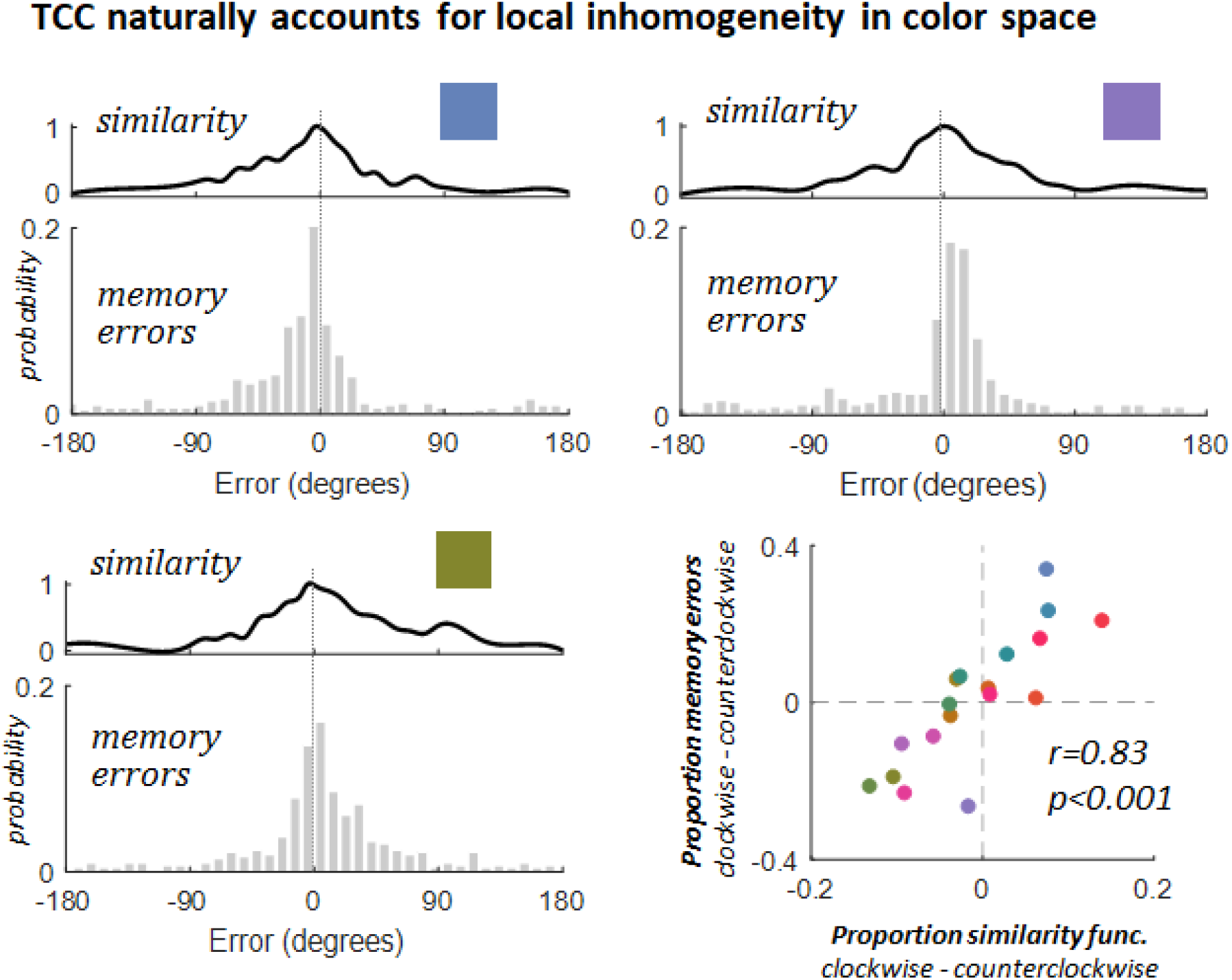
Non-uniformities in memory and similarity for set size data reported in the main text. Many stimulus spaces contain non-uniformities, which may affect subsequent working memory performance. Indeed, Bae et al.^12^ discovered non-uniformities in working memory for color, where responses for targets tend to be more precise for some colors than others and can be biased towards nearby categorical anchors (i.e. red, blue, yellow, etc). While many assume randomizing target colors in working memory should account for potential biases arising from a non-uniform feature space, others have suggested these differences may have broader consequences than previously considered^13,14^. A key advantage of TCC is that by taking into account the psychophysical similarity function, non-uniformities within whatever feature space being probed can be automatically captured if psychophysical similarity data is measured separately from each relevant starting point in the feature space (e.g., Figure 1D). In the current work, we mostly use only a single psychophysical similarity estimate averaged across possible starting points and fit memory data averaged across starting points. However, this is not necessary to the TCC framework, and is only a simplification -- if we wish to fit memory data averaged across all targets, we should use similarity averaged across all targets (or use the particular similarity function relevant to each item on each trial). Here we show that rather than using a psychophysical similarity function that averages over all targets, one can also use similarity specific to each possible target, which differ and have predictable consequences for memory in our set size experiment. For example, the propensity of errors (at set size 1, 3, 6 and 8) in the clockwise vs. counterclockwise direction for a given target color is directly predicted by the similarity function -- even when very similar colors have more similar colors in opposite directions (top row), and this is true across all color bins (bottom right). Thus, using target-specific similarity functions naturally captures potential non-uniformities or biases within a feature space with no change in the TCC framework.

**Extended Data Figure 6.**
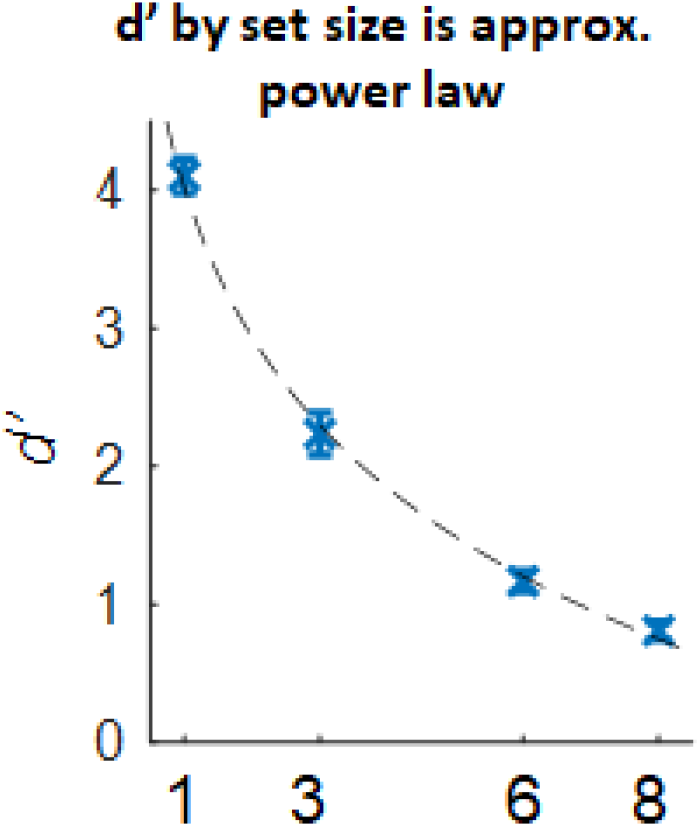
Data from the set size experiment reported in the main text. While memory strength varies according to a variety of different factors, many researchers have been particularly interested in the influence of set size. TCC shows that at a fixed encoding time and with a fixed delay, memory strength (d’) decreases according to a power law as set size changes, broadly consistent with fixed resource theories of memory^10,25^. However, capacity cannot be fixed globally, as the total “capacity” appears to smoothly change with encoding time.

**Extended Data Figure 7.**
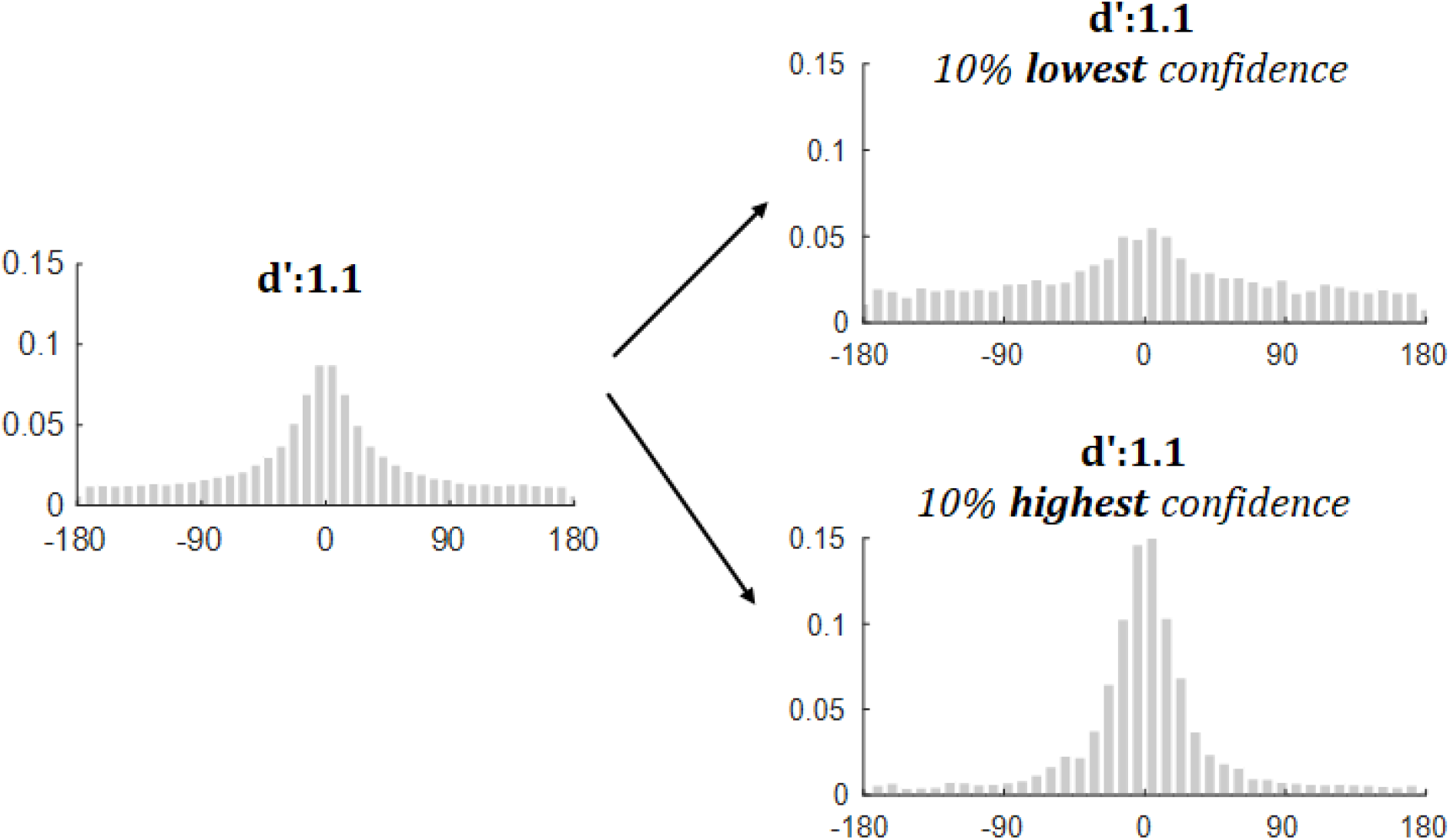
Simulation from TCC illustrating how signal detection can predict variance in representational fidelity as a function of confidence even with a fixed d’ (see also^42^). Some studies used to support variability of information across individual items or trials have done so by using a confidence metric^28^. While variability and confidence are distinct from one another, in a large amount of research they are inextricably linked. An interesting advantage and implication of signal detection-based models is that they naturally predict confidence data^67^. In particular, the strength of the winning memory match signal is used as the measure of memory strength -- and confidence -- in signal detection models of memory. Thus, even with a fixed *d*′ value for all items, TCC naturally predicts varying distributions relative to confidence. This likely explains some of the evidence previously observed in the literature that when distinguishing responses according to confidence, researchers found support for variability in precision among items / trials. Note that this occurs in TCC even though *d*′ is fixed in this simulation -- that is, all trials are generated from a process with the same signal-to-noise ratio. Thus, variability in responses as a function of confidence (or related effects, like improved performance when participants choose their own favorite item to report^23^) are not evidence for variability in *d*′ in TCC, but simply a natural prediction of the underlying signal detection process. Of course, it is possible *d*′ may also vary between items, which remains an open question.

**Extended Data Figure 8.**
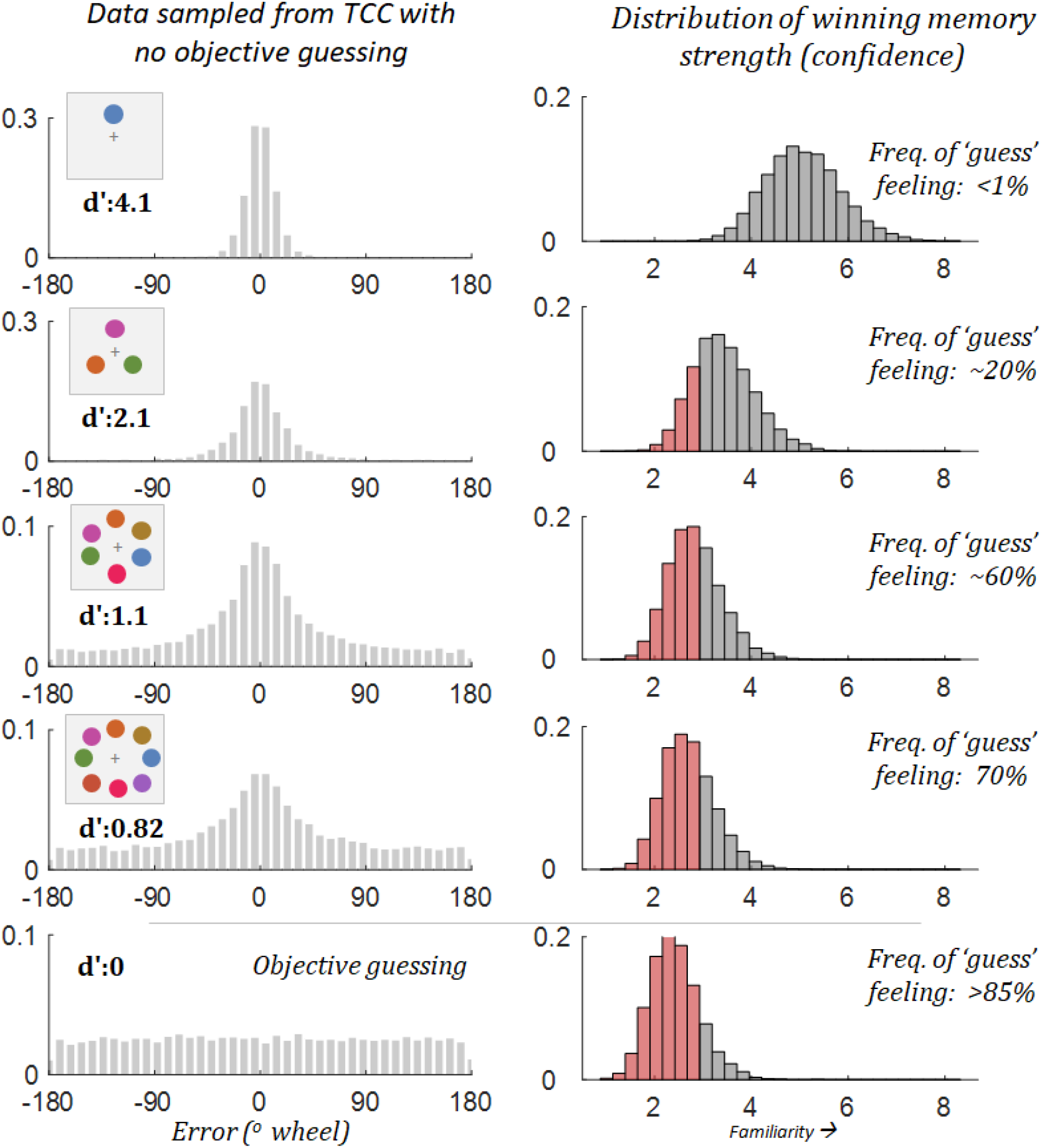
Simulation of confidence in TCC. Participants in a set size 8 working memory experiment often feel like they do not remember an item and are “guessing”, leading to a wide variety of models that predict people know nothing about many items at high set sizes and truly are objectively guessing. However, as noted in Extended Data Figure 7, signal detection naturally accounts for varying confidence, and so can easily account for this subjective feeling of guessing even though in fact, models like TCC predict that people are almost never responding based on no information at all about the item they just saw. In particular, confidence in signal detection is based on the strength of the winning memory signal. Imagine that the subjective feeling of guessing occurs whenever your memory match signal is below some threshold (here, arbitrarily set to 2.75). This would lead to people never feeling like they are guessing at set size 1, and nearly always feeling like they are guessing if they objectively closed their eyes and saw nothing. However, this would also make people feel like they are guessing a large part of the time at set size 6 and 8, even though this data is simulated from TCC -- and the generative process always contains information about all items. This is the key distinction in signal detection models between the subjective feeling of guessing and the claim that people are objectively guessing.

## Supplementary Information

### Supplementary Discussion

#### Measuring psychophysical similarity

The psychophysical similarity function we measure naturally captures two key aspects of how stimuli are perceived: The relationship between the physical stimulus and the psychological representation of that stimulus is rarely linear (e.g., CIELab is a complex transform of light wavelengths), and the similarity between stimuli as a function of distance is additionally non-linear^17,19^. In spaces that are already scaled to be approximately psychophysically uniform (e.g., CIELab), then, only the approximately-exponential fall-off in similarity remains to be modeled; whereas in spaces that are not equalized in advance (e.g., face space), both factors will be measured together, and inhomogeneities may need to be taken into account when modeling memory (e.g., Extended Data Figure 5, fitting each color separately).

In the current manuscript, we present several examples of tasks that naturally capture both of these insights and can be translated to a psychophysical similarity function, including the triad task, the quad task, and a subjective Likert similarity judgment (see Methods). It is important to note that depending on the number of trials, a large number of data points (and many subjects) may be necessary in order to obtain reliable estimates of a given stimulus space in the triad and quad tasks (in the current methods we collected n = 100 participants and pooled across them completely to obtain reliable group estimates). A Likert similarity task may be sufficient to capture this function under some circumstances, like for color in the current study. In such tasks, participants are simply asked to rate the similarity of two items (varying in distance from one another) on a Likert scale from 1 to 7, and these ratings can then be normalized. In color space, we observed this similarity rating task provided a measure of psychophysical similarity that is in close agreement with the results of the quad and triad tasks and requires considerably less data to estimate (Figure 1).

However, it is important to note that depending on the stimulus space, observers may utilize different strategies in such subjective similarity tasks, and that ultimately objective tasks like the quad task may be best to understand the psychophysical similarity function. In particular, to ensure the similarity function is properly measured, is important to ensure that participants provide judgments of the absolute interval between stimuli and not rely on categories or verbal labels, or, in the triad task, that participants not rely on a relational or relative encoding of the two choice items rather than their absolute distance to the target item (that is, the modeling assumes they compare each choice entirely separately to the target item -- not relying on comparing the two choices, say, considering which choice is more clockwise in an orientation task). How best to ensure that participants rely on absolute intervals is represented in a large literature dating to Thurstone^64^ and Torgerson^13^.

Multidimensional stimuli, like color or faces, seem to have general agreement across many methods of measuring psychophysical similarity. However, we expect that collecting the psychophysical similarity measurements will be particularly challenging in single-dimensional stimulus spaces whose true objective distance function is transparent to participants. For example, when asking to judge orientation similarity or location similarity along a circle, participants are likely to be aware that the stimuli are physically manipulated on only a single dimension (angle), and will thus be inclined to report linear similarity judgments along this dimension. Less transparent similarity tasks, like the quad task, may help with this, but it may ultimately be difficult to prevent participants from using this knowledge. How best to deal with this remains a question for future work. For example, it may be possible to instead “back out” the similarity function from memory data, or from alternative tasks (like speeded same-different tasks), or to use speeded similarity tasks to reduce such cognitive strategies. Alternatively, performing multidimensional scaling on the stimuli to create a psychophysically uniform space (as in CIELab for color; for example, in orientation this would “stretch” the space near the cardinals and shrink it near the obliques), could allow relatively simple similarity models. After such scaling, it would be likely that the similarity function beyond the perceptual discrimination limit would be an exponential function, which could allow the parameterization of the similarity function in relatively straightforward terms without the need for complex measurements.

It is important to note that while we emphasize the stability of the similarity function across conditions in the current work, the psychophysical similarity we measure could not possibly be a fixed property of the colors per se, but must be at least partially contextual. For example, if the background color of the display was blue rather than light gray, this would certainly alter the perception of -- and discriminability of -- colors from each other, as would adaptation and many other factors^49,50^, which would necessarily have consequences for memory.

In addition, extremely brief presentations or extremely long presentations that allow verbal coding would be expected to alter this similarity function. It is expected this would result in changes in memory performance as well, in the same way that observed memory biases are altered when discriminability is affected by adaptation or contextual effects^48^. Thus, while we find the similarity function is fixed across a wide range of encoding times, delays and set sizes, there are likely to be conditions which change the underlying perception of the memoranda (e.g., very very short encoding times; different backgrounds) which will necessarily have an effect on memory.

#### “Dissociating” guess rate and precision

In addition to fitting a two parameter model, some previous research has claimed to dissociate these parameters. If a one-parameter model can account for the data, how has previous research so often found dissociations between these parameters?

The majority of these dissociations find that precision (SD) does not change when the ‘guess rate’ (or capacity) does change^7,31^. However, this dissociation is naturally explained by TCC because at low d′ values, ‘guess rate’ can change by a huge amount with SD changing by only a few degrees. For example, over a wide range of guess rates, precision may only vary between SD=21 and SD=24, a difference that is visually indistinguishable and would require extremely high power to detect (e.g., Supplementary Figure 4). As an example, sampling 20 subjects of 100 trials each of data from the TCC at *d*′=1.0 vs. *d*′=0.7 and fitting these data with the 2-parameter mixture model reveals that such an experiment would find p<0.05 for ‘capacity’ greater than 60% of the time but p<0.05 for ‘precision’ approximately 11% of the time, despite both parameters being necessarily linked in the data from TCC. In line with this interpretation, many researchers have now found that with high enough power, previous studies claiming only a change in ‘guess rate’ but not ‘SD’ actually find changes in both, with very small changes in SD present along with large changes in ‘guess rate’^65^. Other dissociations have sometimes been found -- for example, Zhang and Luck^7^ report a manipulation that causes a change in SD but not ‘guess rate’ -- but these dissociations inevitably rely on comparisons across different sets of stimuli with different psychophysical similarity functions (e.g., the Zhang and Luck manipulation adds color noise to the items, making them less distinct), which is perfectly consistent with TCC.

### Supplemental Figures

**Supplementary Figure 1.**
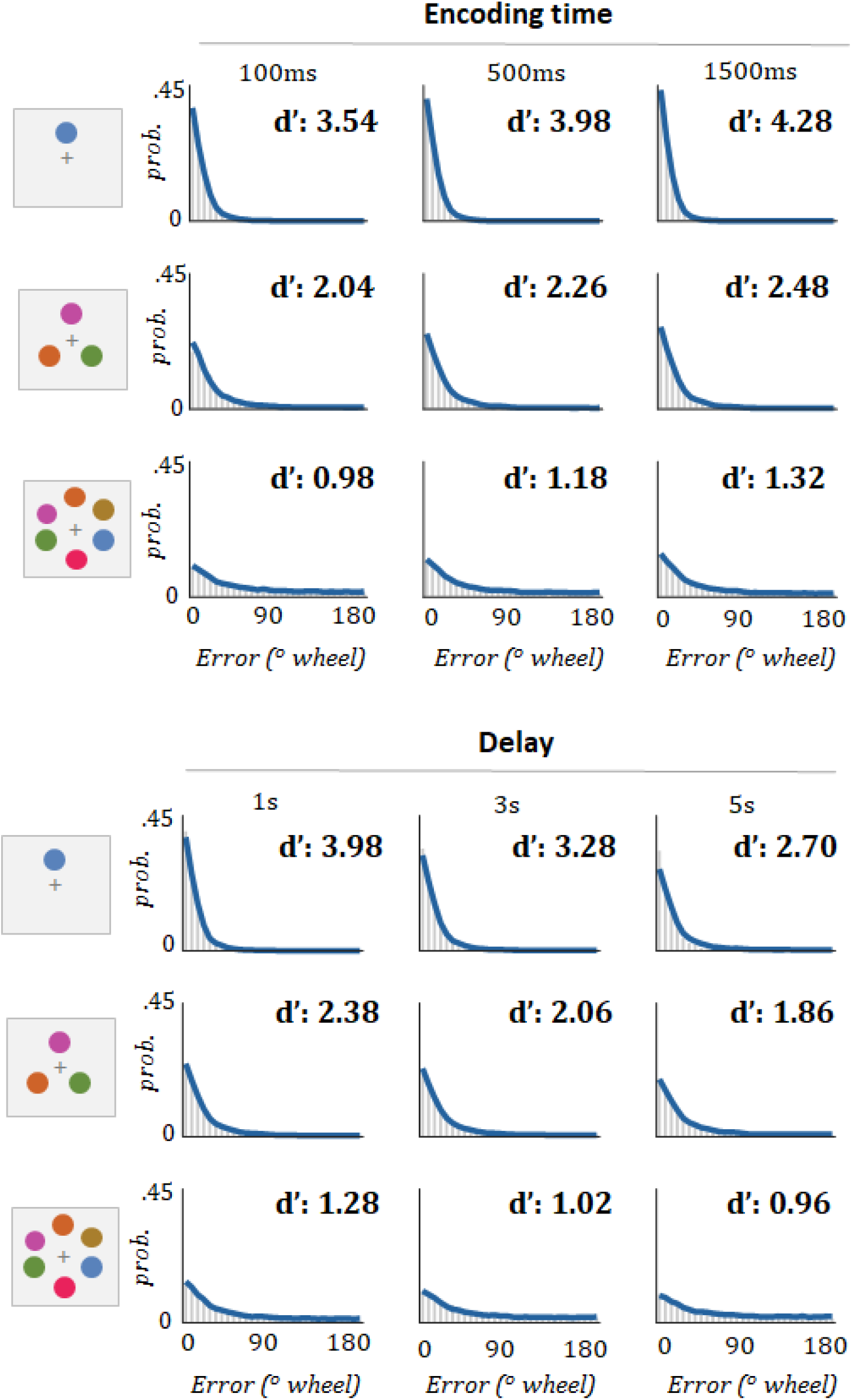
Fits of TCC to the all encoding and delay conditions, including those not plotted in Fig. 3. TCC provides a strong fit at all encoding and delays (see correlations and model comparisons in Fig. 3).

**Supplementary Figure 2.**
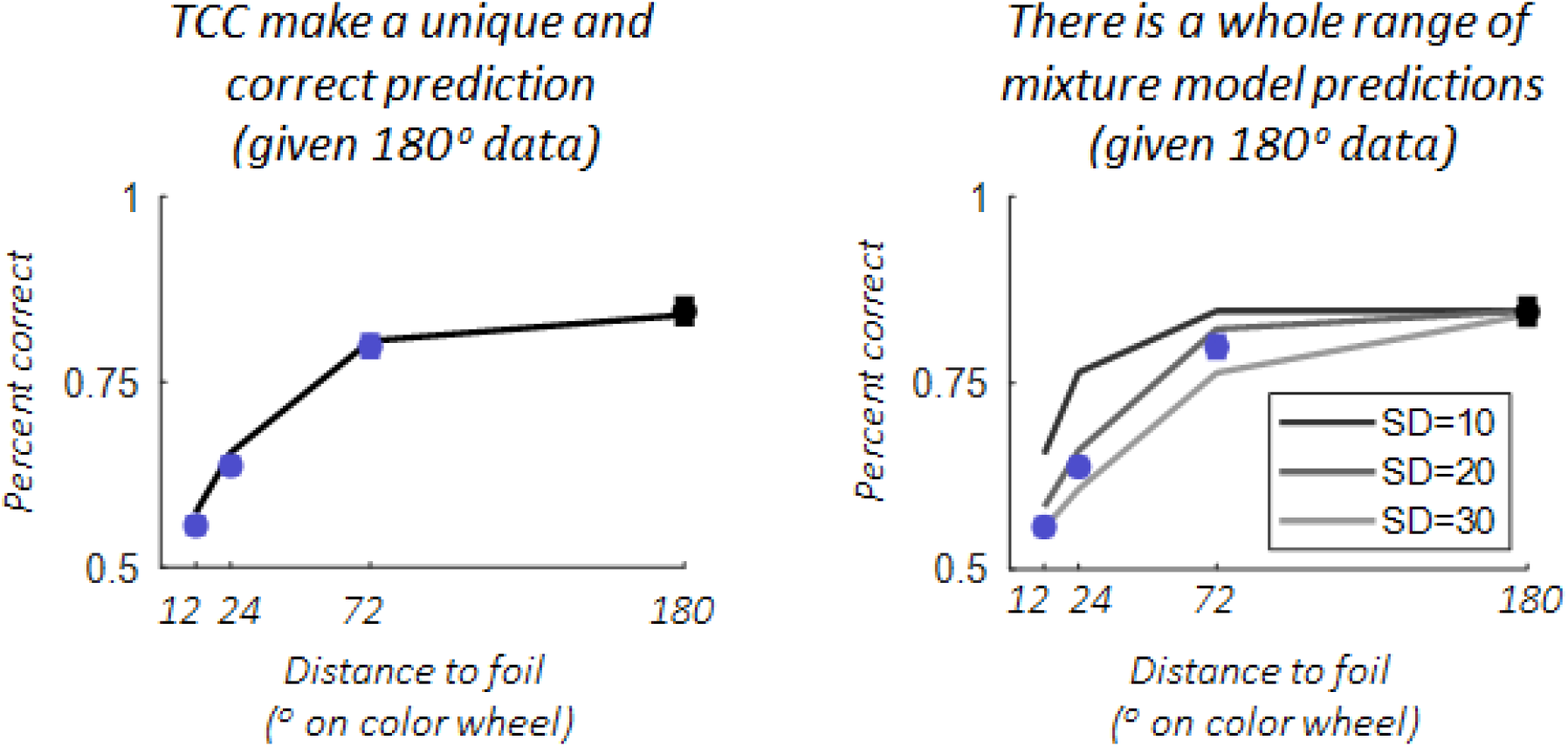
Model predictions vs. data for 2-AFC generalization task reported in the main text. Given 2-AFC performance with maximally distinct 180 degree foils (black dot), TCC makes a unique prediction about exactly how well people should perform on other foils -- with no free parameters. By contrast, using the 180 degree foils to constrain the mixture model allows this model to set the ‘guess rate’, but it leaves the precision of memory unknown. Thus, mixture models, while capable of fitting the data the same as TCC for a certain precision parameter (since ultimately they can predict any distribution TCC can, as they are much more flexible), do not make a unique prediction. Making strong predictions is the most critical test of a model^26^ and can be formalized using a Bayes factor, which provides strong evidence in favor of TCC in this case. Similar logic applies in the experiment taking 180 degree 2-AFC and generalizing to continuous report and other n-AFC conditions.

**Supplementary Figure 3.**
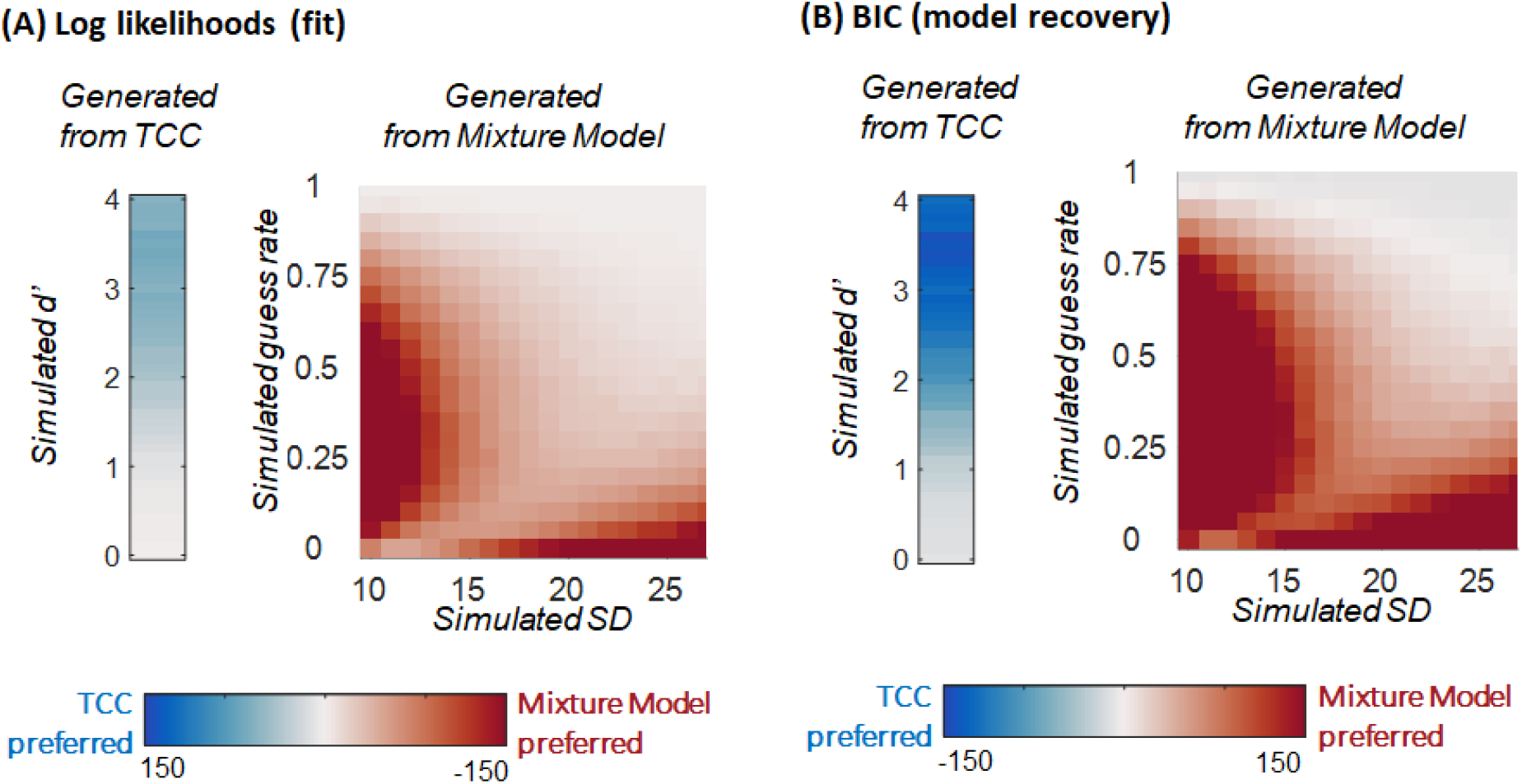
Simulation of mixture model vs. TCC fits. (**A**) We generated data from both TCC (*d*′) and the standard mixture model (precision [SD] and guessing), performing 50 simulations of 2000 trials worth of data each for each of the models (consistent with the amount of group data in the main experiments), and then fit both models to the generated data to see which yielded a higher log-likelihood. With no penalty for complexity -- simply using log likelihood -- for data generated by TCC, the standard mixture model fit all data with a *d*′ < 1 better than TCC itself. Thus, for data generated by TCC, the standard mixture model, being considerably more flexible than TCC in the range of distributions it can fit, fits the data about as well -- and in some cases, better -- than TCC. When fitting data generated by the mixture model, TCC was dispreferred at all values in terms of fit, and strongly dispreferred for huge swaths of potential mixture model parameters. This is because the mixture model can generate a huge variety of distributions that TCC cannot mimic. The same is true, but even more so, for the 3-parameter variable precision model, which can fit an even much larger range of distributions than even the standard 2-parameter mixture model. Only a miniscule part of the distributions predicted by the 3-parameter variable precision model can even be approximated by TCC, and this model can perfectly mimic TCC. (**B**) Same data, with BIC instead of log-likelihood. Taking into account model complexity increases the preference for TCC in TCC-generated data and creates a very slight TCC preference in mixture model data with simulated “guess rates” very near 1.0, where the two models make identical predictions in terms of error (of equal responding to all options); though note the two models make differing predictions about confidence at these values, predicting different ROCs. In general, with this amount of data, BIC appears well-calibrated, accurately recovering the appropriate model in nearly all cases and with a stronger preference for the relevant models where they diverge from each other more.

**Supplementary Figure 4.**
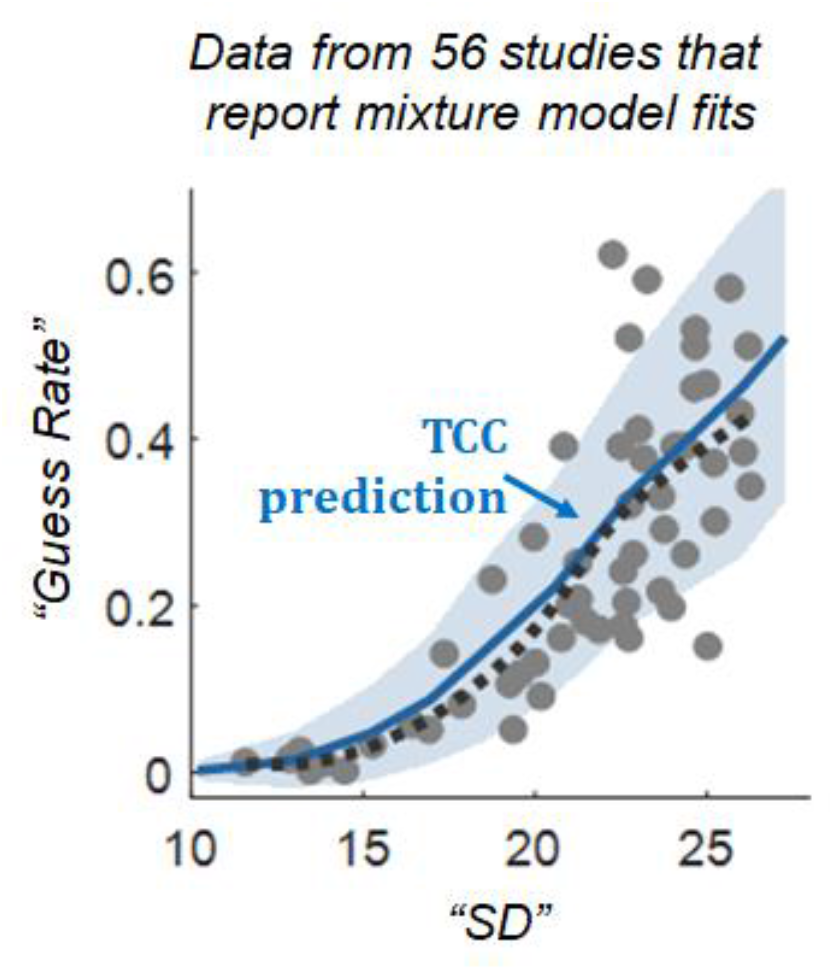
Analysis of previous literature measuring the most widely used model parameters currently used to analyze working memory performance. Gray dots are values reported in papers found in the literature; the dashed black curve is a LOESS (local regression) smoothed version of these points. The solid blue curve reflects the average “guess” and “SD” parameters when fitting the mixture model to data generated by TCC, as a function of the *d*′ of TCC. The blue shading shows 2 standard deviations when each participant has 100 trials/condition. Despite claiming to independently model multiple parameters, this entire diverse set of data points falls near the trade-off between these parameters predicted when fitting data sampled from the TCC with the 2-parameter model -- in other words, one parameter is sufficient to capture much of the data observed in working memory tasks (data that has previously been thought to require at least two -- and often 3 parameters -- to explain). Note that the region in Supplementary Figure 4 TCC predicts is also the only region of Supplementary Figure 3 where the TCC can fit data generated from the mixture model. In addition, note that some of these papers use different color wheels than the one we use to generate the similarity function, and thus some of the deviation from the TCC prediction line -- minor as it is -- is caused by using an “incorrect” TCC prediction (e.g., using a prediction from an incorrect stimulus space).

**Supplementary Figure 5.**
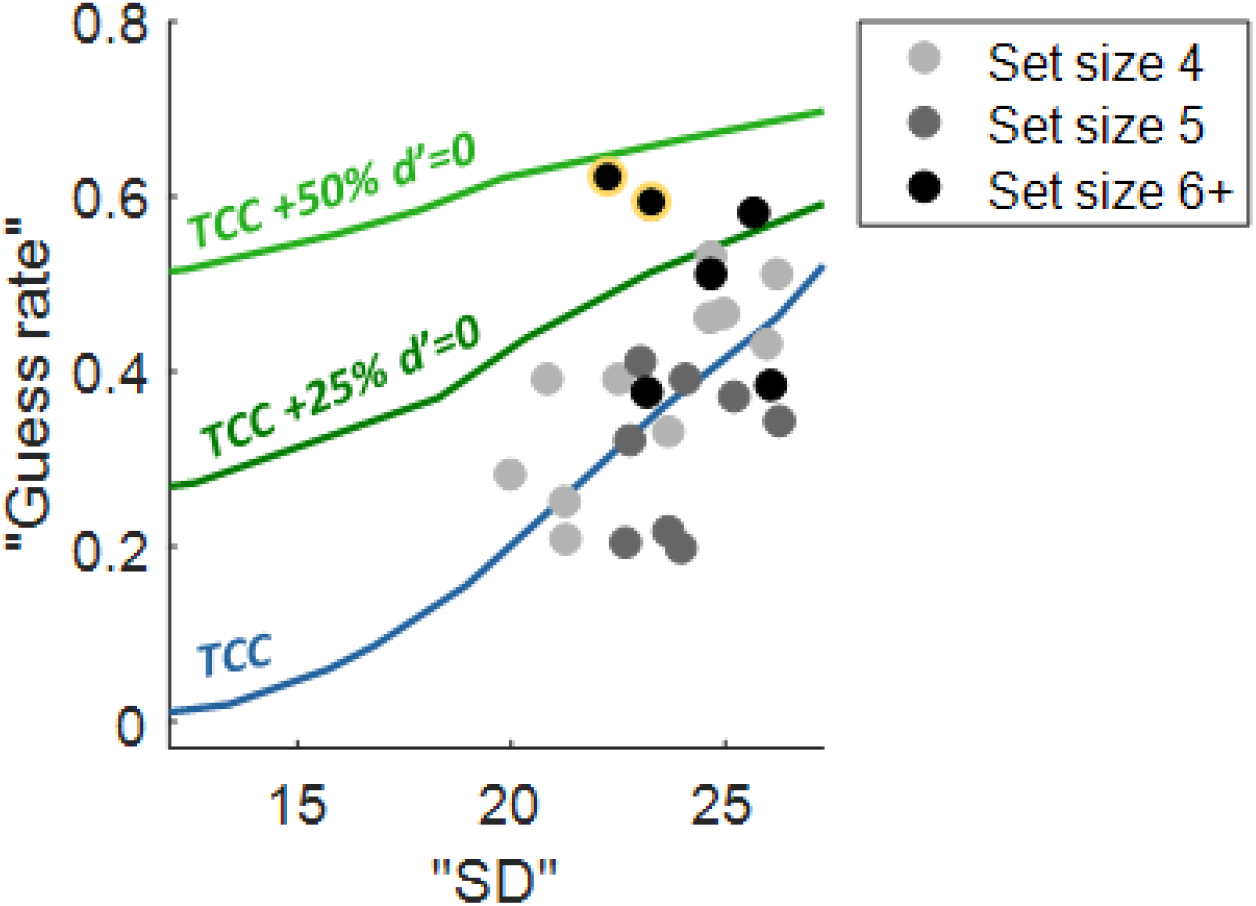
Analysis of previous literature measuring the most widely used model parameters currently used to analyze working memory performance. Existing working memory data from high set sizes (4+) is often claimed to provide evidence for ‘slots’ or for the existence of very low precision items, with these items that are unrepresented or poorly represented giving rise to the long tails of the distribution. By contrast, TCC predicts such long tails with no sense of unrepresented or poorly represented items. Here we show how TCC predicts that mixture model parameters from the standard two parameter mixture model should change as a function of *d*′ in the TCC model. The blue line and all of the data points are the same as Supplementary Figure 4, but with the data points now labeled by set size and only “high” set sizes (>=4) plotted, as these are the points where traditional models claim many items must be unrepresented or extremely poorly represented. Note that the vast majority of the points are better fit by the straightforward TCC model -- which simply assumes all items are equally well represented -- than by models that add some proportion of ‘unrepresented’ items to TCC (plotted in green; note that as expected, these models selectively change the predicted ‘guess rate’ parameter). For a slot model prediction with 3 items represented, nearly 50% of items should be unrepresented at set size 6, and this is clearly incompatible with the previous data as well as the data we report in the main manuscript. In general, the parameters found in the previous literature are perfectly consistent with the basic TCC prediction with no added assumptions about unrepresented items or poorly represented items. Note that the two set size 6 points outlined in yellow come from the original Zhang and Luck^7^ paper that introduced mixture models to this literature and used them to argue for slots. The fact that they are an outlier on this plot may be the reason those authors proposed a model that argues that only ‘guess rate’ but not ‘standard deviation’ changes as a function of set size.

**Supplementary Figure 6.**
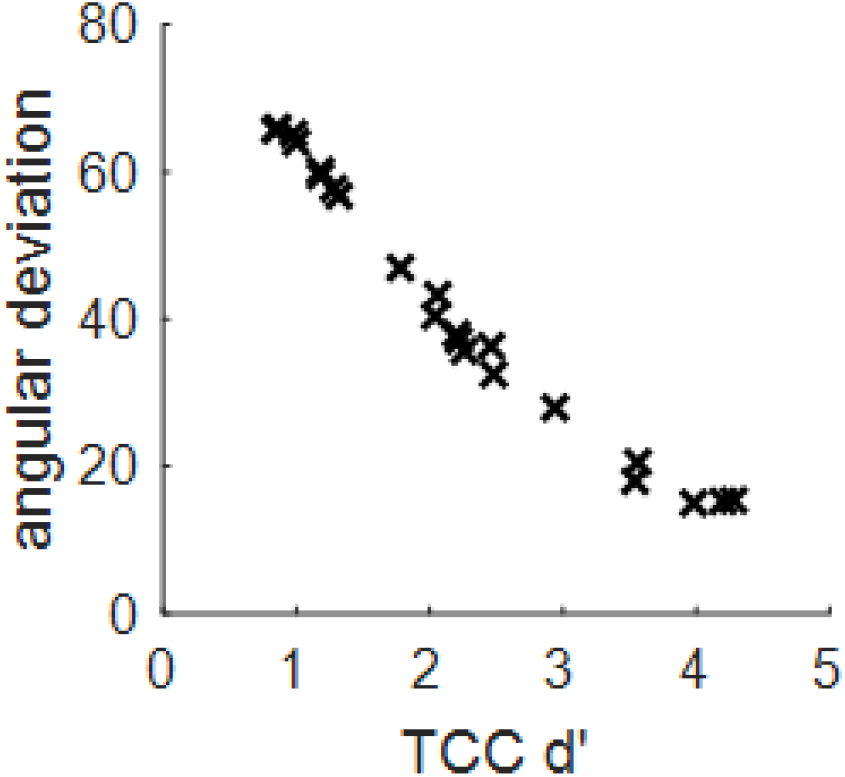
Plot of the best fit TCC d’ vs. the circular standard deviation of the error data (a circular analog of the standard deviation; as computed with MATLAB’s circ_std function) for all 22 datasets from Fig. 3. For data like the current data where there is nearly no location-based confusions (‘swaps’), the simpler analysis of this descriptive statistic (circular standard deviation, or more formally the angular deviation) is linearly related to d’ for d’ less than approximately 3.0, and thus, for data not near ceiling, may be an adequate substitute for fitting the full TCC. This is useful because the circular standard deviation is just a descriptive statistic of the data and thus does not require the collection of similarity data or perceptual confusability data. Note that just as with percent correct -- which is approximately linear with d’ when far from ceiling, but becomes deeply non-linear near ceiling -- the d’ curve begins to bend near ceiling. This is because improving from 95% correct to 99% correct requires a very large change in d’, and similarly, improving your performance in continuous report when it is already very good requires a large change in memory strength. In theory the same should be true near floor, although these 22 datasets do not clearly demonstrate that because there is little data with d’<1.0. However, for data away from ceiling and floor and with little or no ‘swaps’, computing circular standard deviation may be sufficient to summarize data in a framework compatible with TCC.

### Supplementary Tables

**Supplementary Table 1.** TCC’s fit to binned color memory errors (Fig. 3). All correlations are Pearson correlations.

| *Set size experiment* |  |
| --- | --- |
| *Set size 1* | $r=0.998$, $p<0.001$,<br>CI=(0.997, 0.999) |
| *Set size 3* | $r=0.996$, $p<0.001$,<br>CI=(0.993, 0.998) |
| *Set size 6* | $r=0.984$, $p<0.001$,<br>CI=(0.969, 0.991) |
| *Set size 8* | $r=0.976$, $p<0.001$,<br>CI=(0.954, 0.987) |

| *Delay experiment* | *1 sec delay* | *3 sec delay* | *5 sec delay* |
| --- | --- | --- | --- |
| *Set size 1* | $r=0.997$, $p<0.001$,<br>CI=(0.994, 0.998) | $r=0.998$, $p<0.001$,<br>CI=(0.995, 0.999) | $r=0.995$, $p<0.001$,<br>CI=(0.990, 0.997) |
| *Set size 3* | $r=0.993$, $p<0.001$,<br>CI=(0.986, 0.996) | $r=0.992$, $p<0.001$,<br>CI=(0.985, 0.996) | $r=0.994$, $p<0.001$,<br>CI=(0.988, 0.997) |
| *Set size 6* | $r=0.989$, $p<0.001$,<br>CI=(0.979, 0.994) | $r=0.971$, $p<0.001$,<br>CI=(0.946, 0.985) | $r=0.986$, $p<0.001$,<br>CI=(0.973, 0.992) |

| *Encoding time experiment* | *100ms encoding* | *500ms encoding* | *1.5 sec encoding* |
| --- | --- | --- | --- |
| *Set size 1* | $r=0.992$, $p<0.001$,<br>CI=(0.984, 0.996) | $r=0.997$, $p<0.001$,<br>CI=(0.994, 0.998) | $r=0.998$, $p<0.001$,<br>CI=(0.996, 0.999) |
| *Set size 3* | $r=0.971$, $p<0.001$,<br>CI=(0.945, 0.985) | $r=0.991$, $p<0.001$,<br>CI=(0.983, 0.995) | $r=0.995$, $p<0.001$,<br>CI=(0.990, 0.997) |
| *Set size 6* | $r=0.975$, $p<0.001$,<br>CI=(0.952, 0.987) | $r=0.993$, $p<0.001$,<br>CI=(0.987, 0.997) | $r=0.990$, $p<0.001$,<br>CI=(0.981, 0.995) |

**Supplementary Table 2.**
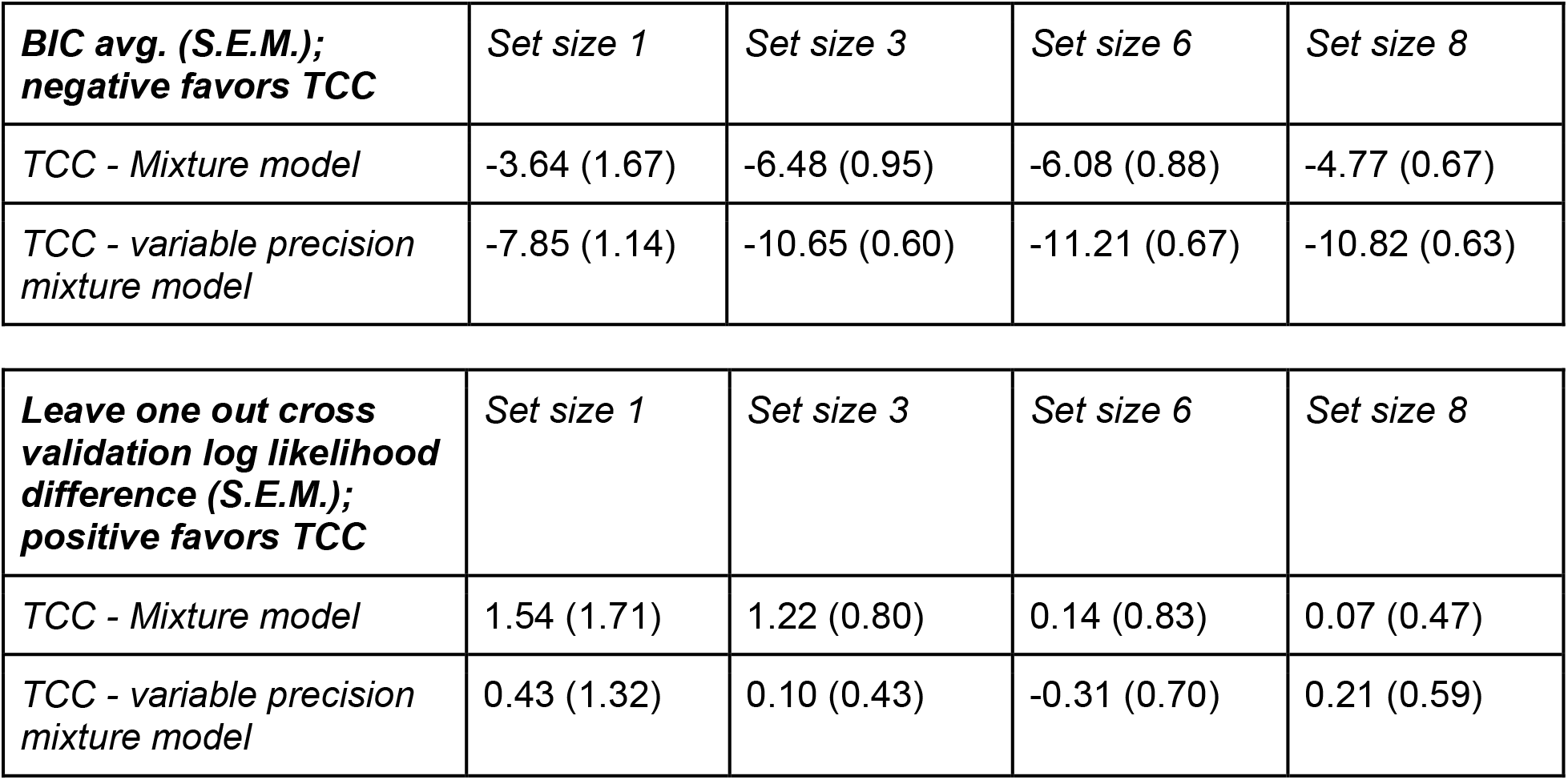
TCC’s fit to color memory data is reliably preferred by model comparison metrics that emphasize simplicity (e.g., BIC) across all set sizes compared to mixture models and variable precision mixture models. It provides a similar fit to these models when using leave-one-out cross validation on log likelihood, as both TCC as well as the two mixture models predict effectively the same distribution of errors when fit with N-1 error points (as N=2000 error datapoints >> the number of parameters for all models). Fitting to the group data rather than individual subjects gives BIC values at set size 1,3,6 and 8 of −24, −56, −26, −25 for TCC vs. standard mixture model (all very strong evidence favoring TCC), and BIC values of −2, −23, −15, −19 for TCC vs. variable precision model (e.g., both models fit set size 1 data well -- the least distinct set size, since there are no long tails -- but all others are very strong evidence in favor of TCC). Note that, as shown in Supplementary Figure 3, model recovery using BIC is well calibrated using this number of trials.

| ***BIC avg. (S.E.M.);***<br>***negative favors TCC*** | *Set size 1* | *Set size 3* | *Set size 6* | *Set size 8* |
| --- | --- | --- | --- | --- |
| *TCC - Mixture model* | -3.64 (1.67) | -6.48 (0.95) | -6.08 (0.88) | -4.77 (0.67) |
| *TCC - variable precision mixture model* | -7.85 (1.14) | -10.65 (0.60) | -11.21 (0.67) | -10.82 (0.63) |

| ***Leave one out cross***<br>***validation log likelihood***<br>***difference (S.E.M.);***<br>***positive favors TCC*** | *Set size 1* | *Set size 3* | *Set size 6* | *Set size 8* |
| --- | --- | --- | --- | --- |
| *TCC - Mixture model* | 1.54 (1.71) | 1.22 (0.80) | 0.14 (0.83) | 0.07 (0.47) |
| *TCC - variable precision mixture model* | 0.43 (1.32) | 0.10 (0.43) | -0.31 (0.70) | 0.21 (0.59) |

**Supplementary Table 3.** TCC applied to face memory. As with colors, TCC is reliably preferred by model comparison metrics that emphasize simplicity (e.g., BIC) across all set sizes compared to mixture models and variable precision mixture models. Also, as with color, it provides a similar fit to these models when using leave-one-out cross validation on log likelihood, as both TCC as well as the two mixture models predict effectively the same distribution of errors when fit with N-1 points (as N >> the number of parameters for all models). Fitting to the group data rather than individual subjects gives BIC values at set size 1 and 3 of −177 and −24 for TCC vs. standard mixture model (all very strong evidence favoring TCC), and BIC values of −53, −10 for TCC vs. variable precision model (all very strong evidence in favor of TCC). Note that, as shown in Supplementary Figure 3, model recovery using BIC is well calibrated using this number of trials.

| ***BIC avg. (S.E.M.);***<br>***negative favors TCC*** | *Set size 1* | *Set size 3* |
| --- | --- | --- |
| *TCC - Mixture model* | -8.1 (0.7) | -5.3 (0.4) |
| *TCC - variable precision*<br>*mixture model* | -11.4 (0.5) | -10.8 (0.3) |

| ***Leave one out cross***<br>***validation log likelihood***<br>***difference (S.E.M.);***<br>***positive favors TCC*** | *Set size 1* | *Set size 3* |
| --- | --- | --- |
| *TCC - Mixture model* | 2.5 (0.46) | 0.51 (0.45) |
| *TCC - variable precision*<br>*mixture model* | 0.87 (0.41) | -0.05 (0.36) |

**Supplementary Table 4.** Data points used in the literature review collected from a total of 14 papers.

| SD | Guess | Paper | Set size | Notes | Digitized |
| --- | --- | --- | --- | --- | --- |
| 13.9 | 0.01 | Zhang & Luck 2008 | SS1 | None | In Paper |
| 19.4 | 0.05 | Zhang & Luck 2008 | SS2 | None | In Paper |
| 21.9 | 0.17 | Zhang & Luck 2008 | SS3 | None | In Paper |
| 22.3 | 0.62 | Zhang & Luck 2008 | SS6 | None | In Paper |
| 20.8 | 0.16 | Zhang & Luck 2008 | SS3 | None | In Paper |
| 23.3 | 0.59 | Zhang & Luck 2008 | SS6 | None | In Paper |
| 22.9 | 0.26 | Zhang & Luck 2009 | SS3 | Retention interval 1 second | In Paper |
| 24.4 | 0.26 | Zhang & Luck 2009 | SS3 | Retention interval 4 seconds | In Paper |
| 24.4 | 0.39 | Zhang & Luck 2009 | SS3 | Retention interval 10 seconds | In Paper |
| 14.5 | 0.001 | Bays, Catalao & Husain 2009 | SS1 | 100ms, Collapsed with swaps | X |
| 19.3 | 0.105 | Bays, Catalao & Husain 2009 | SS2 | 100ms, Collapsed with swaps | X |
| 23.7 | 0.33 | Bays, Catalao & Husain 2009 | SS4 | 100ms, Collapsed with swaps | X |
| 24.7 | 0.51 | Bays, Catalao & Husain 2009 | SS6 | 100ms, Collapsed with swaps | X |
| 13.5 | 0.001 | Bays, Catalao & Husain 2009 | SS1 | 500ms, Collapsed with swaps | X |
| 17.9 | 0.08 | Bays, Catalao & Husain 2009 | SS2 | 500ms, Collapsed with swaps | X |
| 20 | 0.281 | Bays, Catalao & Husain 2009 | SS4 | 500ms, Collapsed with swaps | X |
| 26.1 | 0.383 | Bays, Catalao & Husain 2009 | SS6 | 500ms, Collapsed with swaps | X |
| 13.2 | 0.0245 | Bays, Catalao & Husain 2009 | SS1 | 2000ms, Collapsed with swaps | X |
| 16.45 | 0.0565 | Bays, Catalao & Husain 2009 | SS2 | 2000ms, Collapsed with swaps | X |
| 21.3 | 0.207 | Bays, Catalao & Husain 2009 | SS4 | 2000ms, Collapsed with swaps | X |
| 23.2 | 0.375 | Bays, Catalao & Husain 2009 | SS6 | 2000ms, Collapsed with swaps | X |
| 18.79 | 0.23 | Fougnie, Asplund & Marois, 2010 | SS3 | Single feature | X |
| 25.28 | 0.3 | Fougnie, Asplund & Marois, 2010 | SS3 | Conjunction (w/ orientation) | X |
| 21.5 | 0.18 | Fougnie, Asplund & Marois, 2010 | SS3 | Single feature | X |
| 22.78 | 0.52 | Fougnie, Asplund & Marois, 2010 | SS3 | Conjunction (w/ orientation) | X |
| 24.7 | 0.53 | Zhang & Luck 2011 | SS4 | Low Precision, None | X |
| 26.24 | 0.51 | Zhang & Luck 2011 | SS4 | High Precision, None | X |
| 22.53 | 0.39 | Zhang & Luck 2011 | SS4 | Low Precision, Feedback provided | X |
| 20.87 | 0.39 | Zhang & Luck 2011 | SS4 | High Precision, Feedback provided | X |
| 26 | 0.43 | Zhang & Luck 2011 | SS4 | Low Precision, Payoff provided | X |
| 24.67 | 0.46 | Zhang & Luck 2011 | SS4 | High Precision, Payoff provided | X |
| 11.6 | 0.011 | Fougnie, Suchow & Alvarez 2012 | SS1 | None | In Paper |
| 17.4 | 0.141 | Fougnie, Suchow & Alvarez 2012 | SS3 | None | In Paper |
| 23.7 | 0.216 | Fougnie, Suchow & Alvarez 2012 | SS5 | None | In Paper |
| 21.06 | 0.2 | Brady & Alvarez 2015 | SS1 | None | Lab Data |
| 22.6 | 0.24 | Brady & Alvarez 2015 | SS3 | None | Lab Data |
| 25.7 | 0.58 | Brady & Alvarez 2015 | SS6 | None | Lab Data |
| 22.7 | 0.203 | Fougnie et al 2016 | SS5 | Only included single report condition | In Paper |
| 24 | 0.197 | Fougnie et al 2016 | SS5 | Only included single report condition | In Paper |
| 26.3 | 0.342 | Xie & Zhang, 2016 | SS5 | None | X |
| 23.8 | 0.29 | Suchow, Fougnie, Alvarez 2016 | SS3 | Random report | In Paper |
| 12.9 | 0.015 | Swan, Collins & Wyble, 2016 | SS1 | Pre-surprise | In Paper |
| 15.3 | 0.032 | Swan, Collins & Wyble, 2016 | SS1 | Post-surprise | In Paper |
| 21.3 | 0.25 | Bocincova et al. 2017 | SS4 | None | In Paper |
| 16.9 | 0.05 | Bocincova et al. 2017 | SS2 | None | In Paper |
| 19.6 | 0.117 | Wee et al 2013 | SS3 | Short delay (1 sec) | X |
| 22.6 | 0.174 | Wee et al 2013 | SS3 | Long delay (10 sec) | X |
| 22.8 | 0.32 | Wang et al, 2016 | SS5 | On-probe, 200ms SOA | X |
| 24.1 | 0.39 | Wang et al, 2016 | SS5 | Off-probe, 200ms SOA | X |
| 23.05 | 0.41 | Wang et al, 2016 | SS5 | On-probe, 400ms SOA | X |
| 25.24 | 0.37 | Wang et al, 2016 | SS5 | Off-probe, 400ms SOA | X |
| 20.04 | 0.13 | Fougnie, Asplund & Marois, 2010 | SS3 | Single feature 6 features | X |
| 25.06 | 0.15 | Fougnie, Asplund & Marois, 2010 | SS3 | Conjunction (w/ orientation) 6 features | X |
| 20.21 | 0.09 | Fougnie, Asplund & Marois, 2010 | SS3 | Single feature 6 features | X |
| 22.77 | 0.16 | Fougnie, Asplund & Marois, 2010 | SS3 | Conjunction (w/ orientation) 6 features | X |

